# Genital herpes shedding episodes associate with alterations in the spatial organization and activation of mucosal immune cells

**DOI:** 10.1101/2025.06.24.661157

**Authors:** Finn MacLean, Rachael M. Zemek, Adino Tesfahun Tsegaye, Jessica B. Graham, Jessica L. Swarts, Sarah C. Vick, Nicole B. Potchen, Irene Cruz Talavera, Lakshmi Warrier, Julien Dubrulle, Lena K. Schroeder, Anna Elz, David Sowerby, Ayumi Saito, Katherine K. Thomas, Matthias Mack, Joshua T. Schiffer, R. Scott McClelland, Keith R. Jerome, Bhavna H. Chohan, Kenneth Ngure, Nelly Rwamba Mugo, Evan W. Newell, Jairam R. Lingappa, Jennifer M. Lund, the Kinga Study Team

## Abstract

Herpes Simplex Virus 2 (HSV-2) infection results in variable rates of local viral shedding in anogenital skin. The impact of episodic viral exposures on immune cells in adjacent mucosal tissues, including the genital tract, is unknown. However, any immune responses at this site could impact protective mucosal immunity, tissue homeostasis, and adverse health outcomes. To investigate the impact of HSV-2 on cervicovaginal tract immunity, we applied flow cytometry, immunofluorescent imaging, analysis of soluble immune factors, and spatial transcriptomics to cervicovaginal tissue and blood samples provided by a total of 232 HSV-2 seropositive and seronegative participants, with genital HSV-2 shedding evaluated at the time of biopsy. This unique dataset was used to define and spatially map immune cell subsets and localized gene expression via spatial transcriptomics. HSV-2 seropositivity alone was associated with minimal differences in cervicovaginal and circulating T cell phenotypes. However, the vaginal mucosa during active HSV-2 shedding was associated with alterations in T cell, macrophage, and dendritic cell localization and gene expression consistent with increased immune surveillance, with immune activating and suppressing signals potentially reinforcing mucosal tissue homeostasis.

**Summary:** In context of episodic HSV-2 shedding, immune cells mobilize and co-localize in the vaginal epithelium, expressing cytotoxic and inflammatory genes and immunoregulatory genes that collectively may promote tissue homeostasis in settings of episodic viral shedding to limit damage.

## Introduction

Immunologists have classically divided infection into two discrete categories – acute and chronic. Acute infections, such as influenza virus, norovirus, and pertussis, tend to resolve within days or weeks. In contrast, chronic infections, including Hepatitis C virus, tuberculosis, HIV-1, and CMV, are characterized by infection persisting over months to years, with persistent exposure of the host to pathogen-derived antigen plus inflammation. Studies of chronic infections have revealed adaptations in immune responses to restrict immunopathology in settings of host response to persistent antigen exposure, including the development of T cell exhaustion (Trautmann et al., 2006; Urbani et al., 2006; Wherry et al., 2003). Rapid microbial clearance during acute infection, however, leaves the host with functional immune memory that can be recalled upon subsequent exposure to the same infection. Infection by some notable pathogens of public health significance do not fit this dichotomy but rather exhibit episodic infection, wherein exposure to antigen and/or inflammation is occasional or frequent but not persistent. This includes infection with human herpes viruses including HSV-2.

HSV-2 is a highly prevalent and lifelong infection, with an estimated 545 million people infected worldwide as of 2020 (Harfouche et al., 2025). Following initial infection of the genital skin, HSV-2 ascends through neurites via retrograde transport and establishes latency most commonly in peripheral neuronal ganglia. Cycles of viral latency and reactivation can result in episodic release of virus either in the context of painful genital ulcers, fissures, or more commonly through asymptomatic viral shedding in the genital skin and mucosa (Shin and Iwasaki, 2013).

HSV-2 is a leading cause of genital ulcer disease (GUD), with a recent estimate of 178 million people worldwide suffering from at least one episode of HSV-2-related GUD in 2016, or nearly 5% of the global population of 15-49 year-olds (Looker et al., 2020). This tremendous burden of disease underscores the need to develop improved prevention tools, including therapeutic vaccines, which requires an enhanced understanding of natural immunity to the virus at the sites of viral exposure. Here, we characterize immune cells present in distinct anatomic sites in HSV-2 seropositive and HSV-2 seronegative individuals, as well as a subset of individuals actively shedding HSV-2 virus.

Detailed studies of HSV-2 shedding have revealed that genital shedding rates, though somewhat variable among individuals, are highest during the first year of infection and then stabilize over subsequent years (Phipps et al., 2011; Schiffer et al., 2009; Schiffer and Corey, 2013). Genital sampling at 6-hour intervals for two months demonstrated that shedding occurred a median of 18 times per year in HSV-2 seropositive individuals, although most episodes are cleared in less than 24 hours (Mark et al., 2008). This extremely rapid viral clearance following genital shedding suggests successful tissue immunity. Mathematical modeling indicates that a surprisingly low density of HSV-specific tissue resident T cells is sufficient for repeated effective local elimination of HSV-2 infected cells (Roychoudhury et al., 2020; Schiffer et al., 2010; Schiffer et al., 2009; Schiffer et al., 2013; Schiffer et al., 2018). Furthermore, skin from HSV-2-infected individuals reveals clusters of HSV-specific CD4+ and CD8+ T cells (Peng et al., 2021b; Peng et al., 2012; Zhu et al., 2009; Zhu et al., 2007; Zhu et al., 2013) and regulatory T cells (Treg) (Milman et al., 2016) at sites of HSV-2 lesions and after lesional healing. More recently, we demonstrated transient changes in tissue T cells in HSV-2 lesions, although overall, the number of skin T cells and markers of activation and function were stably maintained within genital skin tissues in response to episodes of HSV-2 reactivation (Dave et al., 2023). Studies of a mouse model of HSV infection have clarified features of this protective tissue-resident memory T cell (T_RM_) response upon HSV challenge (Gebhardt et al., 2009; Iijima and Iwasaki, 2014). Notably, using parabiosis combined with a mouse model of vaginal HSV-2 infection, Iijima and Iwasaki demonstrated that CD4+ T_RM_ establish tissue residency in memory lymphocyte clusters within the vaginal mucosa and were crucial for viral containment upon secondary HSV-2 exposure.

Furthermore, chemokine production by local macrophages maintained these T_RM_ (Iijima and Iwasaki, 2014), thus highlighting the importance of tissue immune cell organization in achieving viral control in the context of shedding. Finally, murine studies and mathematical models demonstrate that a higher density of T_RM_ within reactivation microenvironments predicts more rapid infected cell elimination and lower viral loads (Mackay et al., 2012; Park et al., 2018; Schiffer et al., 2010; Schiffer et al., 2013).

A notable difference between most murine and human studies of anti-HSV-2 immunity is the anatomic site of study. Mouse models of HSV-2 have focused on immune cells in the vagina (Shin and Iwasaki, 2013), whereas many human HSV-2 studies have tracked immune cells within genital skin. CCR5+ CD4+ T cells persist after HSV-2 lesion healing in human genital skin (Zhu et al., 2009), thereby suggesting a potential immune-based mechanism to explain the observed increased risk of HIV-1 acquisition in HSV-2 seropositive individuals (Looker et al., 2017). Assessment of tissue T cells in the genital mucosa of HSV-2 seropositive individuals may provide additional insight, as the genital mucosa including the vagina tract (VT) and ectocervix (CX) are more likely sites of sexual HIV-1 acquisition (Spira et al., 1996). A limited number of human studies of the female genital tract mucosa have demonstrated that HSV-2-specific T cells are present in the CX of HSV-2-seropositive individuals (Koelle et al., 2022; Koelle et al., 2000; Peng et al., 2021a; Posavad et al., 2017; Posavad et al., 2015), though immune cells in the vagina have not been assessed.

We have previously characterized T cells within the human CX and VT and identified distinct activation profiles across genital tract sites compared to cells within the circulation (MacLean et al., 2025; Pattacini et al., 2019; Traxinger et al., 2022; Woodward Davis et al., 2021). Given the remaining gaps in our knowledge of how HSV-2 shedding affects immunity in female genital tract tissues, with implications for sexual transmission of HIV-1 and other sexually transmitted infections, here we comprehensively assess not only cellular profiles associated with HSV-2 infection but also the immune cell localization, organization, and gene expression in the vaginal tissue during episodes of HSV-2 viral shedding.

## Results

### Study population characteristics

Two hundred thirty-two participants (N=135 HSV-2 seronegative and N=97 HSV-2 seropositive) from the Kinga Study (ClinicalTrials.gov ID# NCT03701802) met the criteria to be included in this study. For flow cytometry analyses, 124 HSV-2 seronegative and 85 HSV-2 seropositive individuals provided a VT tissue biopsy, an overlapping but unique set of 124 HSV-2 seronegative and 85 HSV-2 seropositive individuals provided a CX tissue biopsy, and 130 HSV-2 seronegative and 92 HSV-2 seropositive provided a PBMC sample. Soluble immune factors were measured from cervicovaginal tract (CVT) secretions from 114 HSV-2 seronegative and 82 HSV-2 seropositive individuals and in serum samples from 134 HSV-2 seronegative and 97 HSV-2 seropositive individuals.

HSV-2 seropositive individuals were mostly asymptomatic at the time of sample collection, with comparable rates of sores reported at any time in the three months prior to their visit (4.4% HSV-2 seronegative vs 6.2% HSV-2 seropositive) and with no participants self-reported to be taking HSV medication at their enrollment visit. At enrollment, two of 135 seronegative individuals reported a history of GUD in the past 3 months, and of 94 seropositive participants with a response to the question, none (0) reported a history of GUD in the previous 3 months. This low prevalence of GUD is consistent with other studies of individuals in Africa (Celum et al., 2010). Study participants were young (87% under the age of 40), with the HSV-2 seronegative group being relatively younger than the HSV-2 seropositive group (Table 1), likely due to fewer lifetime exposures to HSV-2 in the younger, HSV-2 seronegative group. This cohort was sexually active (median sex acts of 8 per month) in generally monogamous relationships, with 41.7% of all females using hormonal contraceptives. Of note, 32.0% of HSV-2 seropositive individuals (versus 8.1% of HSV-2 seronegative individuals) had a sexual partner living with HIV (and remained HIV-1/2 seronegative throughout this study) (Table 1).

**Table 1.** Demographic information for participants who provided samples for flow cytometry and/or soluble immune factor analyses.

| Demographic Variables |  | HSV-2<br>Seronegative:135 | HSV-2<br>Seropositive:97 |
| --- | --- | --- | --- |
| Age (years) | ≤25 | 60 (44.4%) | 18 (18.6%) |
|  | 25-29 | 30 (22.2%) | 23 (23.7%) |
|  | 30-34 | 19 (14.1%) | 17 (17.5%) |
|  | 35-39 | 17 (12.6%) | 17 (17.5%) |
|  | 40-49 | 7 (5.2%) | 20 (20.6%) |
|  | ≥50 | 2 (1.5%) | 2 (2.1%) |
| Nugent Score | 0-3 | 102 (78.5%) | 59 (65.6%) |
|  | 4-6 | 15 (11.5%) | 12 (12.4%) |
|  | 7-10 | 13 (10.0%) | 19 (21.1%) |
|  | Unknown | 5 | 7 |
| HIV Exposure | Exposed | 11 (8.1%) | 31 (32.0%) |
|  | Unexposed | 124 (91.9%) | 66 (68.0%) |
| Hormonal<br>Contraceptive (HC)<br>Use | Not using HC and had<br>menses in last 35 days | 56 (41.8%) | 41 (43.6%) |
|  | Not using HC and has<br>secondary amenorrhea | 19 (14.2%) | 17 (18.1%) |
|  | Uses HC | 59 (44.0%) | 36 (38.3%) |
|  | Unknown | 1 | 3 |
| Semen Exposure<br>(unprotected sex acts) | Median (IQR) | 8 (3,12) | 8 (3,12) |

To limit the effect of potentially confounding variables on local CVT immunology analyses, we adjusted flow cytometry analysis of the CX and VT tissue samples, as well as soluble immune factor analysis from CVT fluid, for the following variables: age (continuous variable), BV status diagnosed by Nugent score (Nugent et al., 1991) (categories defined in Table 1), HIV exposure (defined as having sexual partner living with HIV), hormonal contraceptive use (categories defined in Table 1), and semen exposure (continuous variable defined as sex acts per month). We adjusted systemic immune analyses, including PBMC flow cytometry analysis and serum soluble immune factor analysis, for age and hormonal contraceptive use to limit the effect of potential confounding variables. We report image analysis results from a subset of participants selected as unexposed to HIV and BV negative (Nugent Score 0-3) (CX: HSV-2 seronegative N=44, HSV-2 seropositive N=19; VT: HSV-2 seronegative N=55, HSV-2 seropositive N=24). The spatial transcriptomic analysis consisted of a smaller subset of individuals described below in the spatial transcriptomics section.

### HSV-2 seropositivity is not associated with a shift in the proportion of T cell subsets or T cell density in the VT or CX

Given the predominance of CD3+ T cells among total CD45+ immune cells in the CVT and blood (Figure 1A) and the potential importance of HSV-2-mediated T cell responses to local viral shedding, we focused our flow cytometry-based analysis on major T cell subsets and their phenotypes (Supplemental Table 1). We separately analyzed the proportion of CD8+ T cells (Figure 1B), conventional CD25-CD127+/- CD4+ T cells (Tconv) (Figure 1C), and CD25+ CD127-Foxp3+ CD4+ regulatory T cells (Treg) (Figure 1D) as a fraction of total T cells across tissue sites comparing HSV-2 seropositive vs HSV-2 seronegative individuals. After adjustments, we found no statistically significant alterations by HSV-2 serology status in the proportions of CD8+ T cells, Tconv, or Treg in any of the three tissue sites.

**Figure 1.**
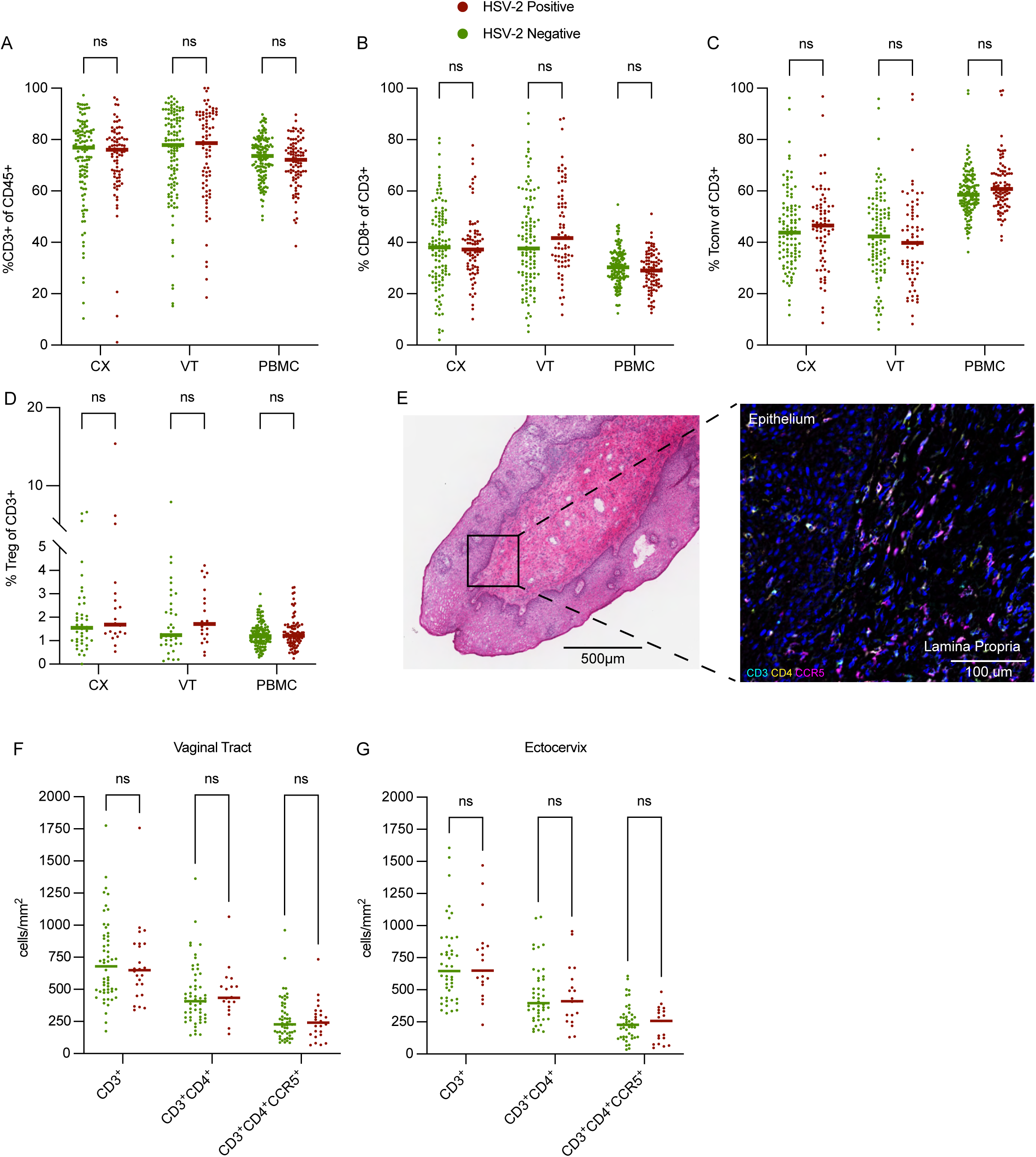
HSV-2 seropositivity does not associate with alterations in total T cells or T cell subsets in the genital mucosa or circulation. (A) Flow cytometry was used to measure the proportion of CD3+ cells among total CD45+ cells in ectocervix (CX) (N HSV-2 seronegative = 122, N HSV-2 seropositive = 83), vaginal tract (VT) (N HSV-2 seronegative = 119, N HSV-2 seropositive = 83), and PBMC samples provided by HSV-2 seropositive and seronegative participants (N HSV-2 seronegative = 130, N HSV-2 seropositive = 92). (B) CD8+ frequency (CX N HSV-2 seronegative = 113, N HSV-2 seropositive = 97; VT N HSV-2 seronegative = 112, N HSV-2 seropositive = 97; PBMC (N HSV-2 seronegative = 130, N HSV-2 seropositive = 92) (C) CD4+ CD25-CD127 +/- Conventional T Cell (Tconv) (CX N HSV-2 seronegative = 103, N HSV-2 seropositive = 74; VT N HSV-2 seronegative = 105, N HSV-2 seropositive = 66; PBMC (N HSV-2 seronegative = 130, N HSV-2 seropositive = 92), and (D) CD4+CD25+CD127-Foxp3+ Regulatory T Cells (Treg) (CX N HSV-2 seronegative = 42, N HSV-2 seropositive = 21; VT N HSV-2 seronegative = 36, N HSV-2 seropositive = 20; PBMC (N HSV-2 seronegative = 130, N HSV-2 seropositive = 92) among total CD3+ T cells in CX, VT, and PBMC from HSV-2 seropositive and seronegative participants. (E) Representative H&E image of VT tissue (left), and representative immunofluorescent-stained VT tissue section from the same sample as shown in the H&E image (right). The box on the H&E image represents the portion of the serial section that is shown in the immunofluorescent image. Quantification of the density of CD3+, CD3+CD4+, and CD3+CD4+CCR5+ cells in (F) VT (N HSV-2 seronegative = 55, N HSV-2 seropositive = 24) and (G) CX (N HSV-2 seronegative = 44, N HSV-2 seropositive = 19) tissue sections comparing HSV-2 seropositive vs HSV-2 seronegative samples. Each dot represents an individual sample, and each bar represents the median. For flow cytometry analysis, comparisons were made using an adjusted rank regression model. PBMC comparisons were adjusted for hormonal contraceptive use and age, and CX and VT comparisons were adjusted for hormonal contraceptive use, bacterial vaginosis via Nugent score, HIV exposure, semen exposure, and age. Image analysis was done on a subset of samples that were unexposed to HIV and BV-negative. Wilcoxon rank sum test was used to compare cellular densities via image analysis without adjustment. “ns” indicates results were nonsignificant with an adjusted P value (for flow cytometry) and an unadjusted P value (for image analysis) greater than 0.05.

In addition to assessing the composition of the T cell compartment across anatomic sites via flow cytometry, we sought to quantify the numbers and location of T cells in the VT and CX through immunofluorescence microscopy. Cryopreserved VT and CX tissue biopsies were sectioned and stained to assess the density and location of CD3+ total T cells as well as CD3+CD4+ T cells (Representative H&E and immunofluorescent image from the serial VT tissue sections shown in Figure 1E). Consistent with our flow cytometry data (Figure 1, A and C), we found no differences in the density of CD3+ or CD3+CD4+ cells in the VT or the CX when assessed as a whole (Figure 1F and 1G) or when analyzing the epithelium and lamina propria of each tissue separately (Supplemental Table 2).

A adverse outcome associated with HSV-2 infection is increased HIV susceptibility (Looker et al., 2017). Previous studies have observed increased potential HIV target cells expressing the HIV coreceptors CCR5+ and CD4+ on T cells in the healing HSV-2 skin lesion (Zhu et al., 2009) and in CVT cytobrushes from HSV-2 seropositive vs HSV-2 seronegative individuals (Rebbapragada et al., 2007; Shannon et al., 2014) without assessing whether this phenomenon is true in the deeper CVT tissue layers of those with HSV-2. We hypothesized that greater abundance of CCR5+ activated CD4+ T cells in the CVT, where sexual exposure to HIV likely occurs, may account for the documented epidemiologic association of increased HIV susceptibility among those HSV-2 seropositive. Therefore, we also assessed CD3+CD4+CCR5+ cell density in the VT and CX and found no differences in either tissue site when assessed as a whole (Figure 1, F and G) or when analyzing the epithelium and lamina propria of each tissue separately (Supplemental Table 2). In total, our flow cytometry analysis and image analyses demonstrate that asymptomatic HSV-2 seropositivity is not significantly associated with alterations in the overall populations of major immune cell subsets in CX, VT, or PBMC samples, nor does it impact T cell and HIV target cell density within the CX or VT tissue layers.

### Vaginal T cells from HSV-2 seropositive individuals displayed altered expression of markers associated with intrinsic and extrinsic immunoregulation

Given the proximity of the VT to probable sites of viral reactivation in the anogenital skin, we hypothesized that HSV-2 seropositivity would correlate with phenotypic alterations of T cell subsets in the VT. In addition to markers used to identify T cells (CD45 and CD3) and the major T cell subsets (CD4, CD8, CD25, CD127, and Foxp3), we also evaluated the expression of Tbet to identify Th1 type Tconv cells (Th1) and CD161 and CCR6 double positivity to identify Th17 type Tconv cells (Th17). To assess if HSV-2 seropositivity was associated with proinflammatory immune responses, we analyzed for the expression of CCR5 as we did by immunohistochemistry, however, this time by flow cytometry, and the activation markers HLA-DR and CD38. To more broadly characterize HSV-2-associated immune alterations beyond markers typically associated with acute inflammation, we assessed the expression of proteins associated with tissue residency (CD69 and CD103), inhibition (CD101 and CTLA-4), dysfunction (CD39), cytotoxicity (Granzyme B), progenitor potential (T Cell Factor 1; TCF-1), exhaustion (PD-1), chemokine trafficking (CXCR3), and memory subsets (CCR7 and CD45RA) following an experimental design previously described (MacLean et al., 2025).

We looked broadly for differences in activation and other phenotypic markers as described above in CD8+ T cells, Tconv, and Treg in the CX and VT (Figure 2A), as well as in Th1 and Th17 cells (Supplemental Table 1). We found a significant increase in the fraction of CD8+ T cells that express CD39 in the VT of HSV-2 seropositive individuals (median HSV-2 seropositive = 9% vs HSV-2 seronegative = 6%; adjusted rank regression beta 2.61, adjusted P value (p_adj_) = 0.0367) (Figure 2B). Additionally, the proportion of CD4+ Tconv cells in the VT that express CD39 was also increased in HSV-2 seropositive individuals (median HSV-2 seropositive = 24% vs HSV-2 seronegative = 14%; adjusted rank regression beta = 4.89; p_adj_ = 0.0844), but did not reach our nominal threshold (p_adj_ <0.05) for statistical significance (Figure 2C). CD39 expression was also observed to be increased, but failed to meet the nominal significance threshold, on CX CD8+ T cells (median HSV-2 seropositive = 9% vs HSV-2 seronegative = 6%; adjusted rank regression beta = 2.16; p_adj_ = 0.0877) with relatively equivalent CD39 expression observed on CX Tconv (median HSV-2 seropositive = 13% vs HSV-2 seronegative = 12%; adjusted rank regression beta = 1.09; p_adj_ = 0.6250) (Figure 2A and Supplemental Table 1), suggesting this phenotypic difference, within the CVT, is more profoundly observed on VT T cells.

**Figure 2.**
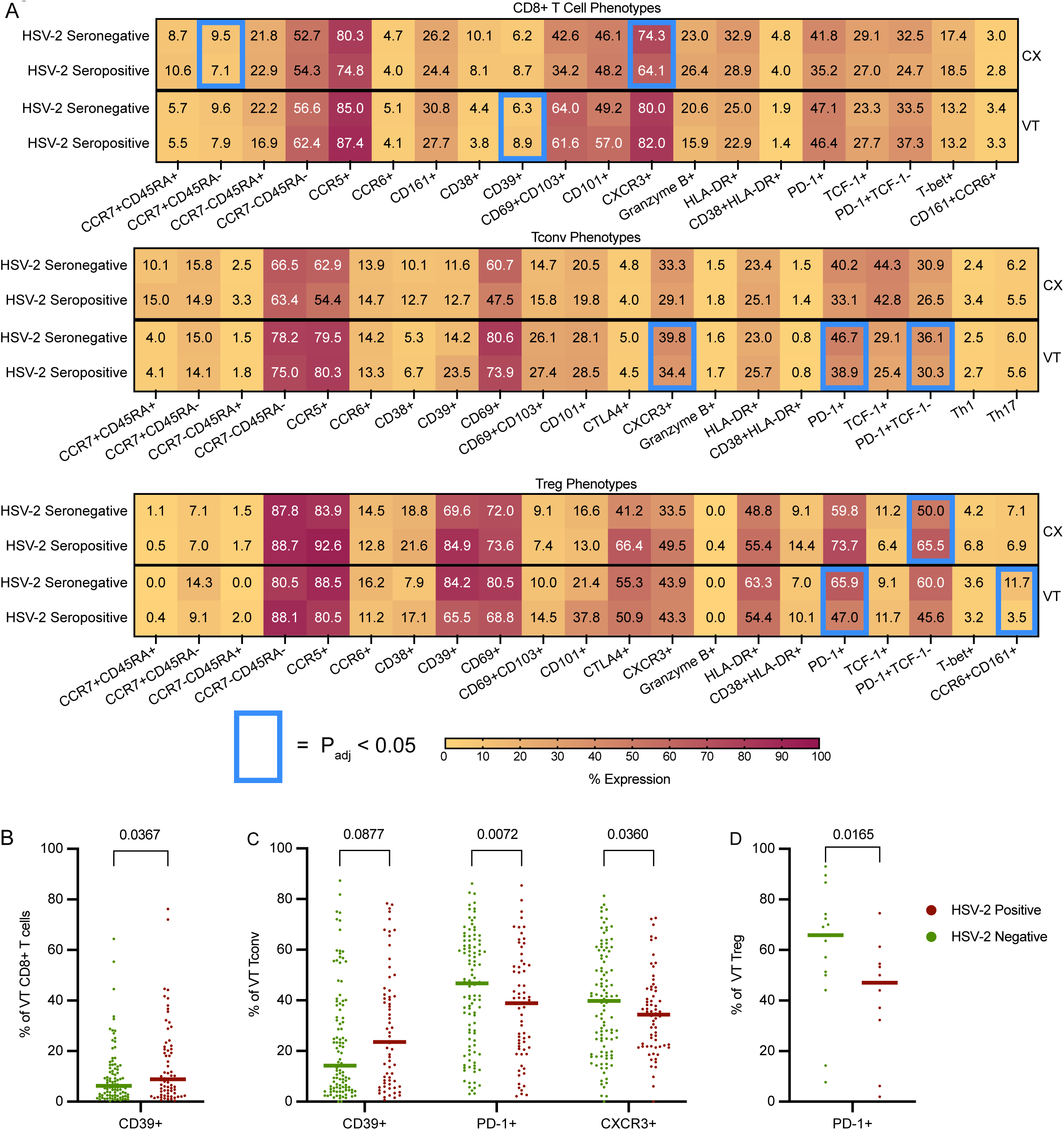
HSV-2 seropositivity drives few phenotypic alterations in the CVT. (A) Heatmap showing frequency of CD8+ T cell, CD4+ CD25-CD127 +/- Conventional T Cell (Tconv), and CD4+CD25+CD127-Foxp3+ Regulatory T Cells (Treg) phenotypes in the ectocervix (CX) and vaginal tract (VT) among HSV-2 seropositive and seronegative individuals. All comparisons were made using a rank regression model that adjusted for hormonal contraceptive use, bacterial vaginosis via Nugent score, HIV exposure, semen exposure, and age. Comparisons on the heatmap that have an adjusted P value < 0.05 are boxed in blue. (CX CD8+ Phenotypes: N HSV-2 Seronegative = 98, N HSV-2 Seropositive = 74; VT CD8+ Phenotypes: N HSV-2 Seronegative = 97, N HSV-2 Seropositive = 69; CX Tconv Phenotypes: N HSV-2 Seronegative = 102, N HSV-2 Seropositive = 73; VT Tconv Phenotypes: N HSV-2 Seronegative = 104, N HSV-2 Seropositive = 22; CX Treg Phenotypes: N HSV-2 Seronegative = 16, N HSV-2 Seropositive = 14; VT Treg Phenotypes: N HSV-2 Seronegative = 15, N HSV-2 Seropositive = 10). (B) Frequency of CD8+ T cells expressing CD39 in the VT of HSV-2 seropositive vs seronegative individuals. (C) Frequency of Tconv expressing CD39, PD-1, or CXCR3 in the VT of HSV-2 seropositive vs seronegative individuals. (D) Frequency of Treg expressing PD-1 in the VT of HSV-2 seropositive vs seronegative participants. Each dot represents an individual sample, and each bar represents the median. Adjusted rank regression P values are displayed. N is the same in panels B, C, and D as described in A.

In addition to altered CD39 expression, we found a significant reduction in the proportion of VT Tconv and Treg that express PD-1 in HSV-2 seropositive individuals (median Tconv HSV-2 seropositive = 39% vs HSV-2 seronegative = 47%; adjusted rank regression beta = -11.48; p_adj_ = 0.0072; median Treg HSV-2 seropositive = 47% vs HSV-2 seronegative = 66%; adjusted rank regression beta = -27.14; p_adj_ = 0.0165) (Figure 2C and 2D) with a reduced effect observed on CX Tconv (median HSV-2 seropositive = 33% vs HSV-2 seronegative = 40%; adjusted rank regression beta = -4.00; p_adj_ = 0.2927) and a marginal, nonsignificant increase observed on Tregs (median HSV-2 seropositive = 74% vs HSV-2 seronegative = 60%; adjusted rank regression beta = 15.98; p_adj_ = 0.1788) (Figure 2A and Supplemental Table 1). While CD39 expression may be associated with impaired effector function and intrinsic regulation, reduced PD-1 expression on VT Tregs may promote greater Treg function and increased T cell-extrinsic immune regulation in the VT. Altered regulatory patterns may be key to shaping tissue memory immune responses to recurrent episodes of viral reactivation. Lastly, we observed a significant decrease in the fraction of Tconv cells that express the chemokine receptor CXCR3 in the VT of HSV-2 seropositive individuals (median HSV-2 seropositive = 34% vs HSV-2 seronegative = 40%; adjusted rank regression beta = -7.90; p_adj_ = 0.0360) (Figure 2C). Decreased frequency of this hallmark receptor of active and infiltrating Tconv may also be related to the regulation of VT immune responses and reduced inflammation. In sum, by applying flow cytometry to CVT biopsies, we identified a limited number of VT T cell phenotypic alterations associated with HSV-2 seropositivity, with fewer significant differences identified in the CX.

### Circulating T cells are primed for trafficking while also displaying signs of regulation in HSV-2 seropositive individuals

In addition to analyzing mucosal immune responses in the CVT in the context of HSV-2 seropositivity, we evaluated whether HSV-2 was associated with signatures of altered phenotypes in circulating T cells (Supplemental Table 1, Supplemental Figure 1). Our analysis revealed that HSV-2 seropositive individuals had an increased proportion of CCR5+ CD8+ T cells (median HSV-2 seropositive = 27% vs HSV-2 seronegative = 21%; adjusted rank regression beta = 3.49; p_adj_ = 0.0278) (Figure 3A) and CCR5+ Th1 cells (median HSV-2 seropositive = 74% vs HSV-2 seronegative = 66%; adjusted rank regression beta = 8.47; p_adj_ = 0.0012) (Figure 3B) in the circulation, which may have the potential to traffic to distal anatomic sites and/or contribute to inflammatory antiviral immune responses. Circulating Tregs from HSV-2 seropositive individuals also more frequently expressed CCR5 (median HSV-2 seropositive = 31% vs HSV-2 seronegative = 27%; adjusted rank regression beta = 3.50; p_adj_ = 0.0265) (Figure 3C), which could contribute to extrinsic regulation of systemic inflammation and/or trafficking to the tissue to provide regulation at the sites of viral reactivation and inflammation.

**Figure 3.**
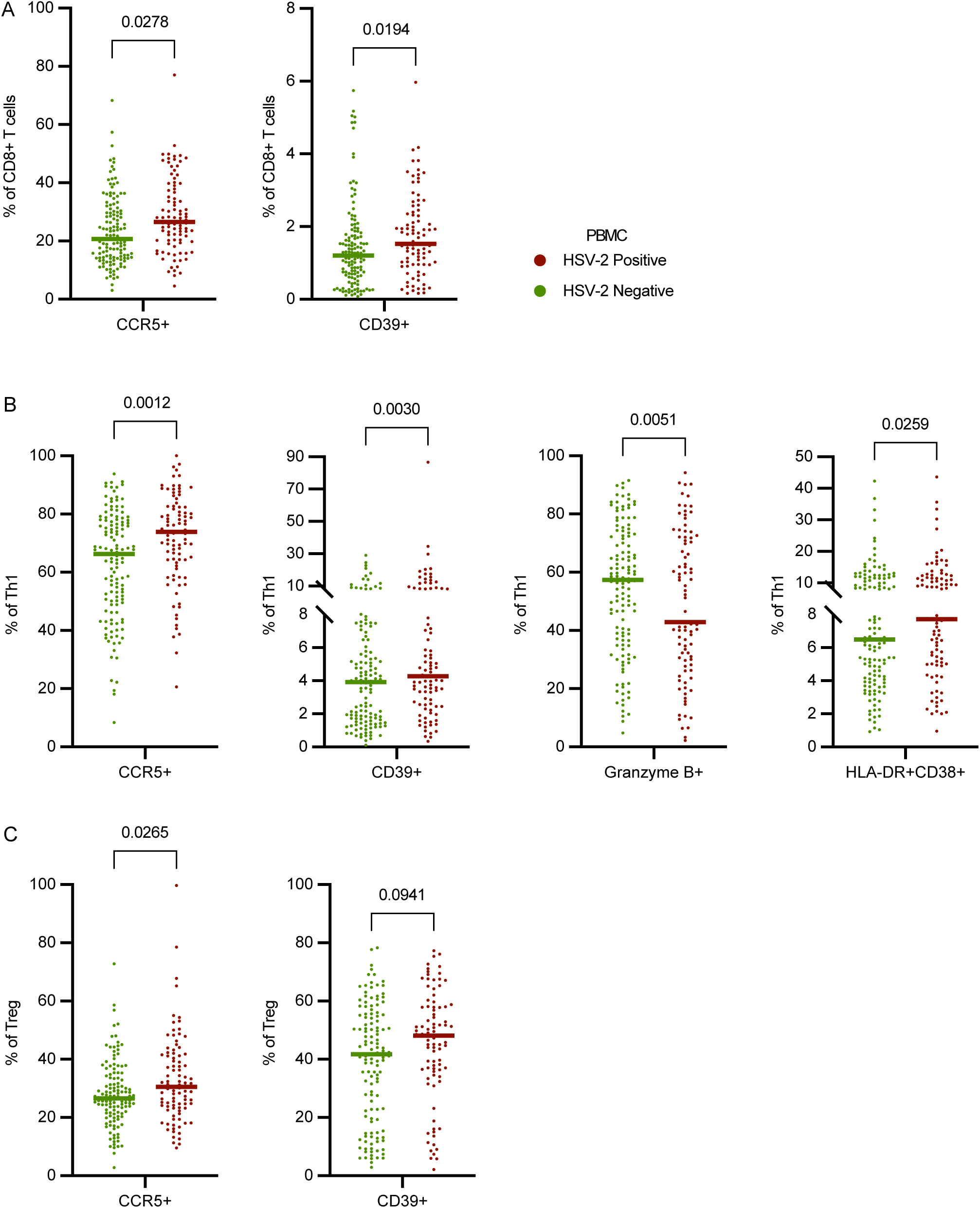
HSV-2 seropositivity is associated with circulating T cell signatures. (A) The frequency of CD8+ T cells in PBMC samples from HSV-2 seropositive vs seronegative participants that express CCR5 or CD39. (B) The frequency of Th1 cells (Tbet+ Tconv) in PBMC samples from HSV-2 seropositive vs seronegative participants that express CCR5, CD39, Granzyme B, or HLA-DR/CD38 double positivity. (C) The frequency of regulatory T cells (Treg) expressing CCR5 or CD39 in PBMC samples from HSV-2 seropositive vs seronegative participants. PBMC comparisons were made using a rank regression model that adjusted for hormonal contraceptive use and age. The adjusted P value is displayed on each graph. Each dot represents an individual data point, and each bar represents the median for that group (N HSV-2 Seronegative = 130, N HSV-2 seropositive = 92).

CD39 was also significantly more frequently expressed by circulating CD8+ T cells (median HSV-2 seropositive = 1.53% vs HSV-2 seronegative = 1.21%; adjusted rank regression beta = 0.32; p_adj_ = 0.0194) (Figure 3A) and Th1 cells (median HSV-2 seropositive = 4.3% vs HSV-2 seronegative = 3.9%; adjusted rank regression beta = 1.23; p_adj_ = 0.0030) (Figure 3B), which may indicate these cells have previously been metabolically active and are subject to intrinsic metabolic regulation. Additionally, there was a trend toward an increase in the fraction of Tregs expressing CD39 (median HSV-2 seropositive = 48% vs HSV-2 seronegative = 42%; adjusted rank regression beta = 4.56; p_adj_ = 0.0941) (Figure 3C). Treg expression of CD39 is associated with greater anti-inflammatory potential and could contribute to Treg-dependent immunoregulation in the context of recurrent antiviral immune responses (Borsellino et al., 2007).

Finally, Th1 from the PBMC samples of HSV-2 seropositive individuals display reduced expression of the cytotoxic effector molecule Granzyme B (median HSV-2 seropositive = 43% vs HSV-2 seronegative = 57%; adjusted rank regression beta = -9.58; p_adj_ = 0.0051), despite having a greater level of activation as measured by CD38+HLA-DR+ expression (median HSV-2 seropositive = 7.7% vs HSV-2 seronegative = 6.5%; adjusted rank regression beta = 1.42; p_adj_ = 0.0259) (Figure 3B). In all, HSV-2 infection may alter systemic T cell phenotypes and drive a systemic signature of differential T cell activation. This signature may reflect a balancing act driven by extrinsic and intrinsic regulatory mechanisms to retain homeostasis and prevent immunopathology while maintaining the potential to readily traffic and respond to viral reactivation in mucosal sites.

### Several soluble immune factors are decreased in serum from HSV-2 seropositive individuals

71 soluble immune factors were analyzed by Eve Technologies Human Cytokine Array/Chemokine Array from the serum and CVT fluid (Softcup®) of HSV-2 seropositive and seronegative individuals to assess the cytokine and chemokine milieu in circulation and the genital mucosa. For soluble factors with any values falling out of the linear range of the assay, these values were imputed (see Methods), and the resulting data analyzed as a continuous variable. If >20% of data were imputed (either above or below the linear range) then the data was analyzed as a dichotomous variable of detected vs not detected (see Methods). In the serum, 53 cytokines/chemokines were quantified as a continuous outcome (Supplemental Figure 2A; Supplemental Table 3), and in the CVT fluid, 61 cytokines/chemokines were quantified as a continuous outcome (Supplemental Figure 3A; Supplemental Table 3). In the serum, 16 factors were analyzed as dichotomous outcomes (Supplemental Figure 2B; Supplemental Table 3), with 2 serum soluble immune factors excluded for being greater than the detectable range for more than 20% of samples and therefore without variation when treated as detectable vs not detectable. Ten factors from CVT fluid were analyzed as dichotomous outcomes (Supplemental Figure 3B; Supplemental Table 3). These analyses identified significant differences in 9 serum factors: PDGF-AA, MIP-1β, Fractalkine, MCP-3, VEGF-A, TNF-ɑ, TNF-β, IL-13, IL-4 (all reduced), and no significant (p_adj_ <0.05) alterations when comparing HSV-2 seropositive vs seronegative CVT fluid samples.

### Spatial transcriptomics identifies tissue and immune cell subsets within VT tissue

While T cells are a dominant population in terms of frequency among CD45+ cells in the cervix and vagina (Figure 1A), we wished to more deeply profile various immune cells subsets within the genital tissues, as well as spatially map their position within the tissue layers. Thus, to assess immune alterations associated with HSV-2 seropositivity as well as active HSV-2 shedding, we used the 10x Genomics® Xenium platform to analyze the location and transcriptional profile of immune cells in VT tissue sections from 5 HSV-2 seropositive and 6 HSV-2 seronegative Kinga Study participants (Table 2). We chose to focus our spatial transcriptomics analysis on vaginal tissue, as this is the anatomic region within the genital mucosa where we identified differences in T cell phenotypes in HSV-2 seropositive individuals (Figure 2A). We also reasoned that HSV-2 shedding from the anogenital skin was more likely to access vaginal tissue due to spatial proximity. Thus, to analyze VT sections from those included in this study, we applied 10x Xenium v1 Spatial gene expression using the Human Multi-Tissue and Cancer panel, consisting of 377 genes. Cell segmentation was performed using Proseg (Jones et al., 2025), which uses cell morphologies identified from DAPI-stained nuclei and the spatial distribution of transcripts to determine cell boundaries. Low-quality cells (<10 probe counts and <5 features) were removed, leaving a total of 142,903 cells. From the panel, we identified four epithelial layers, fibroblasts, endothelial cells, and immune cell types (Figure 4, A and B), which mapped to the anatomically expected areas and structures (Figure 4C, Supplemental Figure 4A). The four epithelial clusters are mapped to the inner layer (stratum basalis), two suprabasal layers, and an outer layer (stratum corneum).

**Figure 4.**
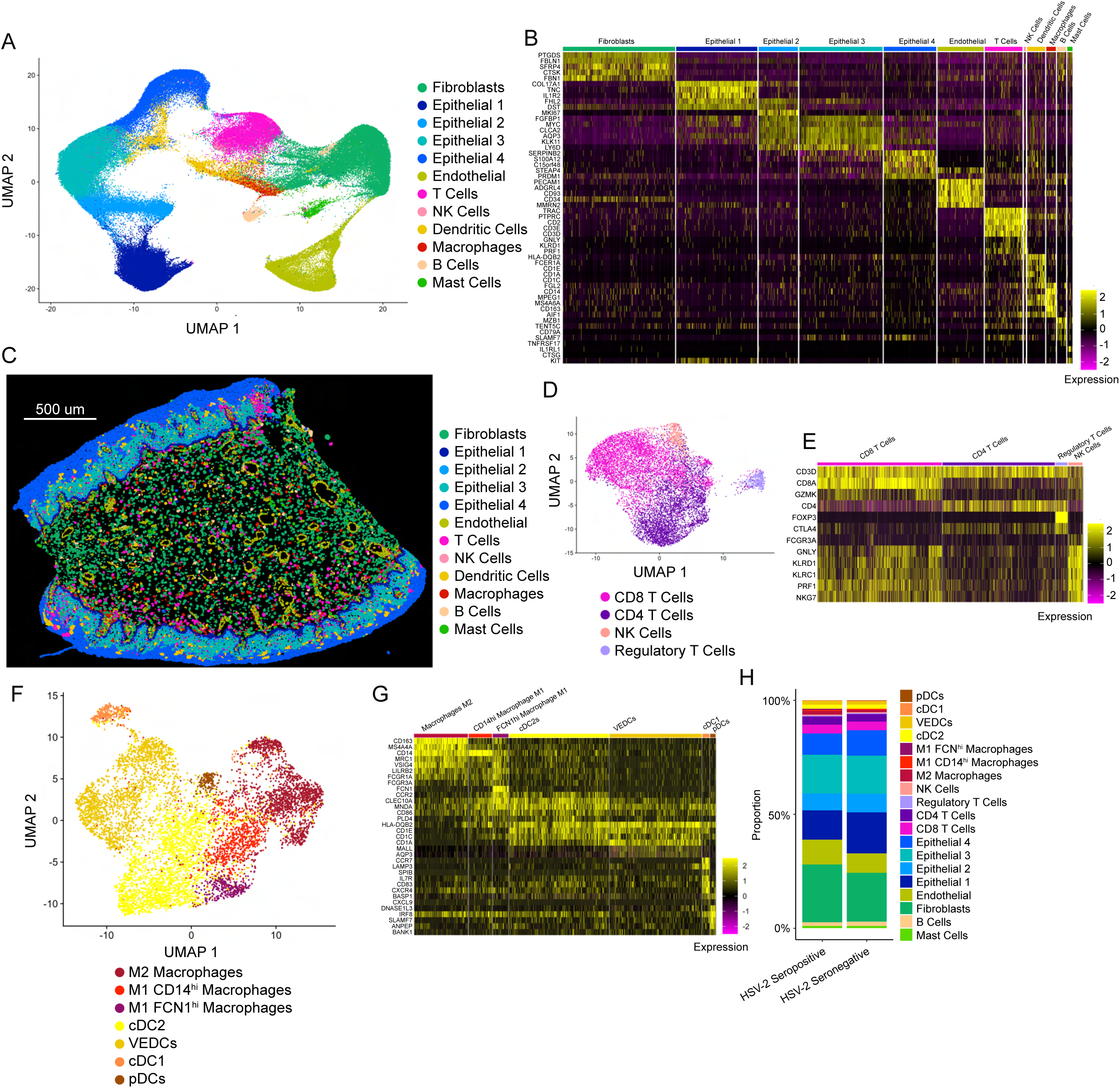
Spatial Transcriptomics identifies cell types and location in the tissue. (A) Annotated UMAP showing the clustering of cells pooled from all samples analyzed on the Xenium platform. (B) Heatmap showing distinguishing transcripts used to identify cell types. (C) The visualization of cells on a representative tissue section. (D) UMAP used to distinguish T cell subsets and NK cells from the broader T cell cluster. (E) Heatmap showing transcripts used to distinguish subsets in (D). (F) UMAP used to distinguish innate immune cell subsets from the dendritic cell and macrophage clusters in (A). (G) Heatmap showing transcripts used to distinguish subsets in (F). (H) Comparisons of cell types present in the tissue between N= 5 HSV-2 seropositive and N=6 HSV-2 seronegative individuals.

**Table 2.** Spatial transcriptomics clinical and demographic data.

| Demographic Variables |  | HSV-2<br>Seronegative:6 | HSV-2<br>Seropositive:5 |
| --- | --- | --- | --- |
| Age (years) | ≤25 | 4 | 1 |
|  | 25-29 | 0 | 0 |
|  | 30-34 | 0 | 1 |
|  | 35-39 | 2 | 2 |
|  | 40-49 | 0 | 1 |
|  | ≥50 | 0 | 0 |
| Nugent Score | 0-3 | 5 | 3 |
|  | 4-6 | 1 | 0 |
|  | 7-10 | 0 | 0 |
|  | Unknown | 0 | 2 |
| HIV Exposure | Exposed | 0 | 0 |
|  | Unexposed | 6 | 5 |
| Hormonal Contraceptive<br>(HC) Use | Not using HC and had<br>menses in last 35 days | 6 | 2 |
|  | Not using HC and has<br>secondary amenorrhea | 0 | 0 |
|  | Uses HC | 0 | 3 |
|  | Unknown | 0 | 0 |
| Semen Exposure<br>(unprotected vaginal sex<br>per month) | Range | 3-12 | 2-18 |
|  | Not reporting sexual<br>activity | 1 | 1 |
| Anogenital Swab HSV-2<br>PCR Result | Positive | 0 | 3 |
|  | Negative | 0 | 2 |
|  | Not done | 6 | 0 |

The T cell/NK cell cluster was subclustered to identify CD8+, CD4+, Treg, and NK cells (Figure 4, D and E). The macrophage and dendritic cell (DC) clusters were subclustered to identify subtypes including M1-like macrophages (CD14hi and FCN1hi populations), M2-like macrophages (CD163+MRC1+), conventional DC (cDC)-2 populations (CD1c+CD1a+), cDC1 (CCR7+LAMP3+), and plasmacytoid DCs (pDC) (IRF8hi SLAMF7+ANPEP+) (Figure 4, F and G). The two cDC2 populations distinguish themselves as a more cDC2-like population (CD1c and CLEC10A high), and the other similar to a vaginal epithelial DC (VEDC) phenotype (CD1a high) which resides in the outer stratum corneum layer.

After identifying cellular populations in the vaginal tissue sections, we compared the cellular composition in samples from HSV-2 seropositive and HSV-2 seronegative individuals. Similar to what we detected by flow cytometry for T cell categories (Figure 1), we found no significant difference in the proportion of any cell type between HSV-2 seropositive and HSV-2 seronegative samples (Figure 4H, Supplemental Figure 4B). In sum, through the use of spatial transcriptomics, we were able to classify a comprehensive range of tissue types, stratifying the mucosal layers, submucosa and endothelial cells in vaginal tissue. Furthermore, we were able to define subsets of immune cell types, thereby allowing for comparison of the proportions of individual cell types.

### VT samples from participants with active HSV-2 shedding contain more immune cells with an inflammatory phenotype

While we were able to identify select T cell phenotypes in the genital mucosa and circulation that associate with HSV-2 seropositivity, we predicted that active viral shedding may have a more pronounced effect on the local genital mucosal immune cell localization. HSV-2 seropositive participants provided an anogenital swab at the time of tissue biopsy collection.

Three participants had detectable HSV-2 virus, confirming they were actively shedding HSV-2 virus at the time of biopsy collection (Shed+), and 2 HSV-2 seropositive participants had a negative anogenital PCR sample and served as the comparator group (Shed-). For HSV-2 seropositive vs seronegative spatial transcriptomics analysis, all 5 seropositive participants, regardless of shedding status, were included. Given the small N included in our spatial transcriptomics analysis, no statistical adjustments were made for potentially confounding variables. Of note, each participant included in this analysis was unexposed to HIV, and there were no participants who were known to be BV+, though some clinical data are missing from individual participants. One HSV-2 Shed+ participant reported having sores on their genital area in the previous 3 months but none at the time of biopsy collection, whereas the other 2 Shed+ and the 2 Shed-reported no sores on their genital area at their visit or at any time in the last three months.

Because we hypothesized that active HSV-2 anogenital shedding may have distinct effects on vaginal immune cells, we next examined samples from HSV-2 seropositive individuals, comparing participants who were actively shedding HSV-2 virus to those with a negative anogenital swab PCR result. Within the T cell compartment, Shed+ samples showed a shift from CD8+ to CD4+ dominant, with a nonsignificant increase in Treg cells (Figure 5, A and B). In terms of cellular profiles, Shed+ individuals had a lower percentage of CD4+ and CD8+ T cells that expressed CD27, CCR2, and GZMK, and a higher percentage that expressed CD69, CLCA2, and CTLA4 genes. CD8+ T cells also displayed increased expression of GZMB in Shed+ samples, suggesting a shift to an activated, cytotoxic phenotype. A greater percentage of Tregs expressed IL2RA and CTLA4 in Shed+ samples, required for Treg survival and suppressive function. NK cells were also more activated in the Shed+ samples, with a greater percentage expressing GZMA, GZMB, and KLRD1 (Figure 5C, Supplemental Figure 5A).

**Figure 5.**
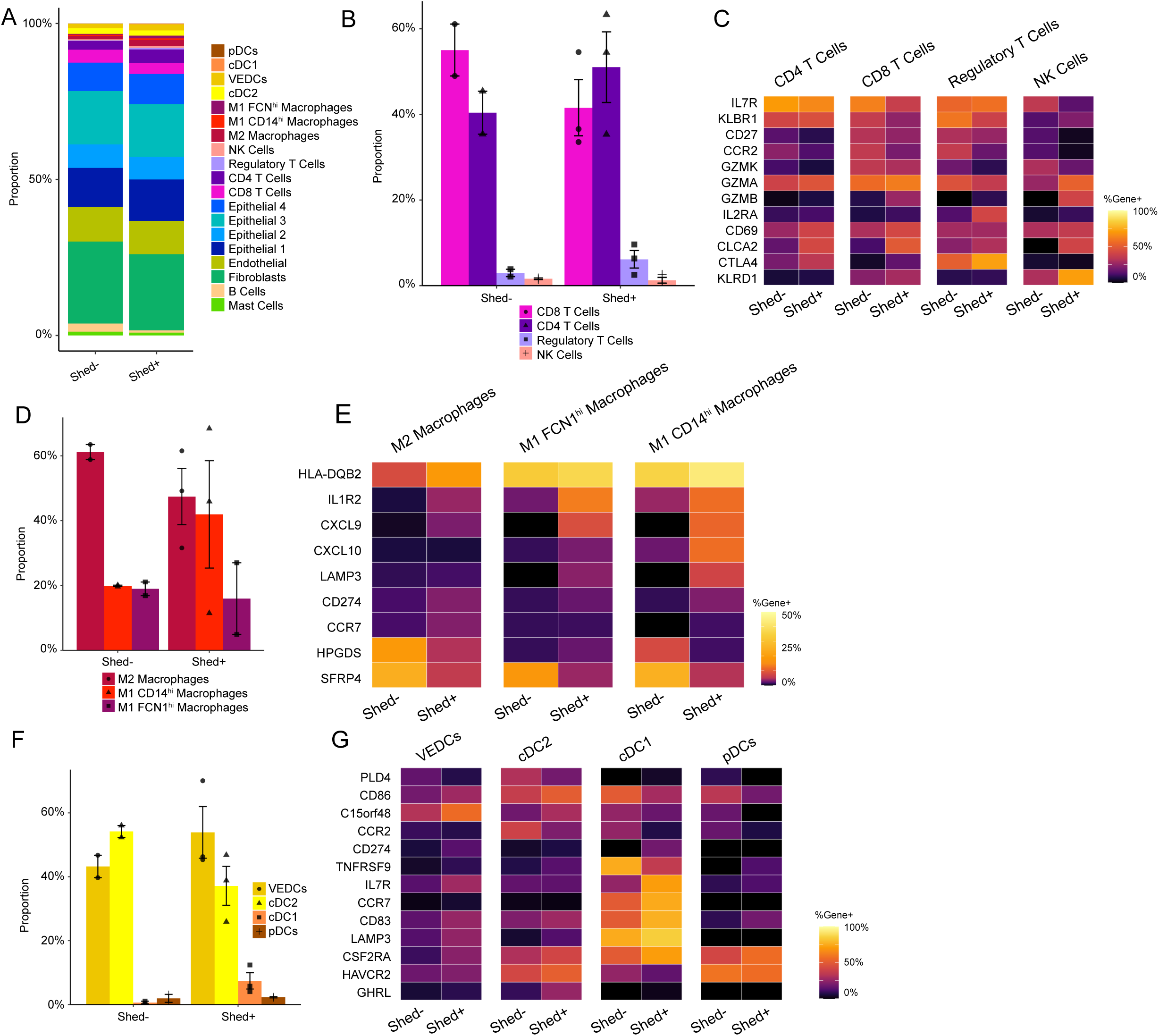
HSV-2 viral shedding associates with inflammatory immune response in the vagina. (A) Proportion of cell subsets in HSV-2 seropositive, anogenital swab HSV-2 PCR negative (Shed-) (N=2) vs HSV-2 seropositive, anogenital swab HSV-2 PCR positive (Shed+) (N=3). (B) Comparison of T cell subsets and NK cells as the frequency of total T and NK cells. (C) Heatmap showing relative gene expression in T and NK cell subsets in Shed-vs Shed+ samples. (D) Comparison of macrophage subsets as the frequency of total cells in the macrophage cluster. (E) Heatmap showing relative gene expression in macrophage subsets in Shed- vs Shed+ samples. (F) Comparison of dendritic cell subsets as the frequency of total cells in the dendritic cell cluster. (E) Heatmap showing relative gene expression in dendritic cell subsets in Shed- vs Shed+ samples.

We also assessed the effect of HSV-2 shedding on macrophage subsets. Shed-samples had predominantly M2 macrophages, while Shed+ samples had a greater proportion of M1 macrophages, particularly the CD14hi M1 population (Figure 5D). These M2 macrophages expressed HPGDS and SFRP4 in the Shed-samples, indicative of a wound-healing or anti-inflammatory phenotype. In Shed+ samples, a greater percentage of M1 macrophages, particularly CD14hi M1 macrophages, expressed IL1R2, CXCL9, CXCL10, and LAMP3, genes known to be upregulated in inflammatory settings (Gonzalez et al., 2015; Tuersun et al., 2024; Yu et al., 2024) (Figure 5E, Supplemental Figure 5B).

Similarly, in the DC compartment, Shed+ samples showed a shift from cDC2 (CD1c+ CLEC10A+) to cDC1 (CCR7+LAMP3+CD83+) cells, with significantly more cDC1 cells in Shed+ samples (Figure 5F). Notably, a greater percentage of cDC1 expressed CD83 and LAMP3 in individuals with active shedding, possibly indicating increased maturity and activation in response to viral stimuli. Further, increased CCR7 expression in cDC1 of Shed+ samples may indicate increased DC maturation and ability to migrate toward the draining LN (Figure 5G, Supplemental Figure 5C). The cDC1 cells also expressed more IL7R and CSF2RA, suggesting they may be migratory or monocyte-derived. In Shed+ samples, more CD1a-high Vaginal Epithelial Dendritic Cells (VEDCs) expressed C15orf48, potentially indicative of an inflammatory response (Clayton et al., 2021). In contrast, Shed-samples had more DCs with CCR2 expression, suggestive of an immature phenotype (Figure 5G, Supplemental Figure 4C). Altogether, analysis of immune cell subsets and gene expression by different types of immune cells in the vagina of HSV-2 seropositive individuals indicates that active HSV-2 shedding is associated with modest shifts in composition of the immune cell compartment, as well as changes toward more inflammatory gene expression.

### Active HSV-2 shedding is associated with macrophage polarization and the recruitment of CD4+ T cells into the epithelium

Given the alteration in immune cells and genes associated with cell migration or recruitment, we hypothesized the location of immune cells between the mucosa and submucosa may be altered during active viral shedding. To explore the spatial distribution of immune cells, the BuildNicheAssay function in the Seurat package was adapted to distinguish between the lamina propria and epithelium (Figure 6, A and B). Not only did cell types differ in their location between the two niches, but also their distance from the basal epithelial layer, with FCN1^hi^ M1 macrophages closest to the basal layer within the lamina propria, CD14^hi^ M1 macrophages venturing into the epithelial layer, and VEDCs farthest out into the epithelium. T cell populations were found in both the lamina propria and epithelium, although CD4+ T cells on average ventured the furthest distance into the epithelium (Figure 6C). Comparison of the proportion of cells in each niche revealed that Shed-samples had a higher proportion of T cells in the lamina propria, while Shed+ samples had more T cells in the epithelium and more macrophages in the lamina propria (Figure 6, D-G). This suggests that HSV-2 shedding is associated with differences in the composition of the immune cell compartment within the vaginal tissue layers.

**Figure 6.**
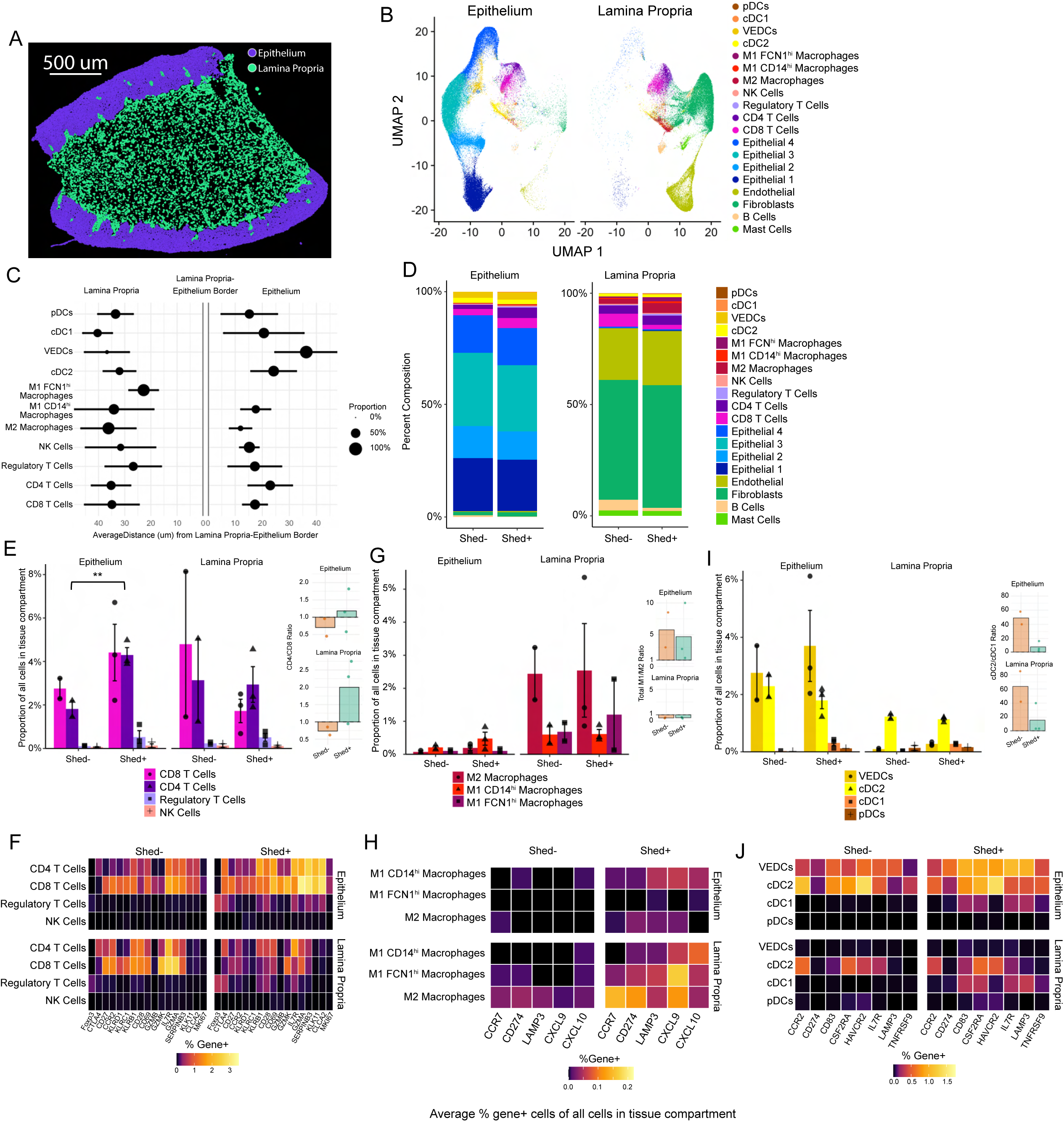
Vaginal inflammatory immune response is enhanced in the epithelium during HSV-2 shedding. (A) BuildNicheAssay function in Seruat was used to stratify the epithelium and lamina propria layers, and (B) clustering cells in each layer confirmed differences in cell populations in each layer. (C) The average distance and distribution of each cell type from the thin basal epithelium cell layer were measured. (D) The proportion of cell types in the lamina propria and epithelium in Shed- vs Shed+ samples was measured. The proportion of T cell subsets and NK cells in Shed- vs Shed+ ( N Shed- = 2, N Shed+ =3) (E) and the heatmap comparing gene expression in T and NK cells in Shed- vs Shed+ (F). The proportion of macrophage subsets in Shed- vs Shed+ (G) and the heatmap comparing gene expression in macrophage subsets in Shed- vs Shed+ (H). The proportion of dendritic cell subsets in Shed- vs Shed+ (I) and the heatmap comparing gene expression in dendritic cell subsets in Shed- vs Shed+ (J).

When further comparing cells within each niche, we found significantly more CD4+ T cells within the epithelium in Shed+ samples compared to Shed-, but no significant difference in the lamina propria (Figure 6E). This was confirmed by calculating the density of T cells (cells per μm2), where the density of CD4 T cells was increased in the epithelium in Shed+ samples, though results were non-significant (Supplemental Figure 6A). Immunofluorescent staining confirmed an increase in the CD3+CD4+ density in the epithelium relative to total CD3+ density in the vaginal tissue. While this difference was not statistically significant (Supplemental Figure 6B), it was a confirmation of our transcript-level findings. Comparing the CD4:CD8 T cell ratio, T cells in Shed-samples were predominantly CD8+ in both niches, while CD4+ cells were the dominant T cell population in both niches in Shed+ samples, particularly in the lamina propria, where the average ratio was 2:1, though this difference was nonsignificant (Figure 6E). This increase in overall CD4+ T cells suggests CD4+ T cells may be recruited from the blood into the tissue in Shed+ samples, maintaining a CD4+ T cell population in the lamina propria as they migrate and increase in number in the epithelium. While the CD8+ T cells make up a greater proportion in the lamina propria in Shed-, this relationship flips in Shed+, where the proportion of CD8 T cells in the epithelium is increased and the proportion in the lamina propria is decreased, consistent with CD8 T cell recruitment from the lamina propria to the epithelium during shedding (Figure 6E). In terms of gene expression (Supplemental Figure 6C), Shed-samples were characterized by more CD4+ and CD8+ T cells in the lamina propria expressing CD27, CCR2, KLRB1, and IL7R, consistent with a resting memory phenotype (Schiott et al., 2004). In contrast, this population is diminished in the Shed+ samples, which have more CD28, CD69, GZMA, KLK11, and CLCA2 expression in CD4+ and CD8+ T cells in the epithelium (Figure 6F), suggestive of cytotoxic activation after interaction with antigen presenting cells. The CD8 T cells in the epithelium of Shed+ samples express the genes for granzymes GZMB+, GZMK+, and GZMA+, effector molecules involved in cytotoxicity of virally infected cells (Figure 6F). CD4 T cells appear to move from the lamina propria in Shed-samples into the epithelial layer in Shed+ samples, while also changing gene expression pattern as seen by the expression of KLRB1+ memory cells from the lamina propria in Shed- to the epithelium in Shed+, where they have loss of CD27, suggestive that these memory T cells are migrating and becoming activated. Additionally, there is increased expression of CTLA4+ on T cells in the epithelium of Shed+ samples, consistent with a regulatory mechanism after immune activation (Figure 6F). This change in gene expression pattern by T cells within the spatial niches, with more inflammatory T cells residing in the lamina propria in Shed-individuals and more cytotoxic T cells and activated CD4+ T cells shifting to the epithelial niche in Shed+, may reflect changes in pathogen-associated molecular pattern (PAMP) exposure within the vaginal lumen and vaginal epithelium that could thereby activate innate immune cells and induce changes in cytokine and chemokine expression.

We next explored the distribution and gene expression characteristics of innate, antigen-presenting cell subsets in the lamina propria and epithelium. Within the macrophage subsets, there was no significant change in the proportions (Figure 6G) or density (Supplemental Figure 6D) of macrophages in the lamina propria regardless of HSV-2 shedding status. There was also no change in the M1:M2 ratio between Shed- and Shed+, although the location influenced this ratio, with M2 macrophages dominant in the lamina propria, and M1 macrophages dominant in the epithelium. Inflammatory genes, including CCR7, CD274, LAMP3, CXCL9, and CXCL10, were increased in macrophages within the lamina propria from Shed+ samples compared to Shed-(Figure 6H). Shed+ samples also had more LAMP3, CXCL9, and CXCL10+ macrophages in the epithelial layer, which corresponded to CD14hi M1 macrophages (Figure 6H). The CD14hi M1 were the main source of CXCL10+ and CXCL9+ macrophages in the epithelium, which was the macrophage subtype that was not only in both the lamina propria and epithelium but also went furthest into the epithelial layers (Figure 6C). Finally, we identified an inverse correlation between macrophage CXCL10/CXCL9 expression and distance from the basal epithelium, particularly in the epithelium in Shed+ samples (CXCL10 r= -0.68, FDR= 6.6^-44^, CXCL9 r =-0.52, FDR=4.9^-35^), with most highly expressing cells within 20µm (Supplemental Figure 6E and F). Taken together, this is suggestive that macrophages, particularly the activated CD14+ M1 subset, may be a source of CXCL10 and CXCL9 chemokines in Shed+ samples, which could in turn attract immune cells such as T cells towards the basal epithelium.

Lastly, within the DC subtypes, Shed+ samples had a small yet distinct cDC1 population in both the lamina propria and epithelial layers, which was not present in Shed-samples (Figure 6I). In addition to LAMP3 and CCR7, this population expressed CD83, CSF2RA, and IL7R, suggestive of an inflammatory migratory population (Figure 6J). There was also a slight decrease in the cDC2 population in Shed+ samples, resulting in a decrease in the cDC2:cDC1 ratio (Figure 6I), though there was no difference in total DC density across experimental groups (Supplemental Figure 6G). The cDC2 population showed a shift from CCR2+ in Shed- to LAMP3+ in Shed + (Figure 6J), indicative of maturation, typically after antigen capture.

Although there was no change in the number of VEDCs associated with active viral shedding, they showed the strongest shift in gene expression, with an increase in cells in the epithelium expressing genes associated with interferon stimulation, such as CD274, HAVCR2, and TNFRSF9, known to be upregulated during active viral shedding (Figure 6J). Altogether, we show that in the context of active HSV-2 shedding, there are changes in the composition of the immune cells in both the lamina propria and the epithelium, and increased expression of activation and inflammatory genes within the epithelial layer in Shed+ samples. This is consistent with mobilization of cells, possibly mediated at least in part through CXCL9/CXCL10 (Supplemental Figure 6E and F), that are involved in immune surveillance and effector function to an anatomic site that may be exposed to viral shedding (the vaginal lumen).

### Inflammatory macrophages may draw T cells toward the epithelium, where they interact with DCs during active HSV-2 shedding

Finally, we sought to identify cellular interactions within different tissue regions to better understand the coordinated networks of the immune response to active viral shedding. To look for cell interactions, distances between cells were calculated, with a distance of 20 μm used as a cutoff for cells that may be directly interacting with one another (see Methods). Using this metric, cells could be classified as near (<20 μm) or far (>20 μm) from a target cell of interest and then queried the frequency of interactions between T cells and antigen presenting cells (macrophages and DCs) by tissue compartment (Figure 7, A and B, Supplemental Figure 7A-C). We first looked at which cells the T cell subsets were co-localizing with and found that most interactions with macrophages occurred in the lamina propria (Figure 7C, Supplemental Figure 7, D and E), with most interactions at the edge of the basal epithelium in Shed+ samples (visual representation, Figure 7A). T cells interacting with DCs were located mostly in the epithelial layer (Figure 7C, Supplemental Figure 7, F and G); however, in Shed-samples, most interactions occurred near the basal epithelium, including in the lamina propria, while in Shed+ samples most of the interactions were throughout the epithelium (visual representation, Figure 7B). In Shed- samples, CD8 and CD4 T cells mainly interacted with cDC2 or M2 macrophages in the lamina propria and cDC2 or VEDCs in the epithelium, with the most abundant interaction between cDC2 and CD8 T cells (Figure 7C). In Shed+ samples, CD4 T cells interacted with cDC1 and FCN1+ M1 macrophages in the lamina propria and cDC1 in the epithelium, CD4, CD8 and regulatory T cells interacted with CD14 M1 macrophages in the epithelium, and both CD4 and CD8 T cells had more interactions with cDC1 and VEDCs in the epithelium (Figure 7C).

**Figure 7.**
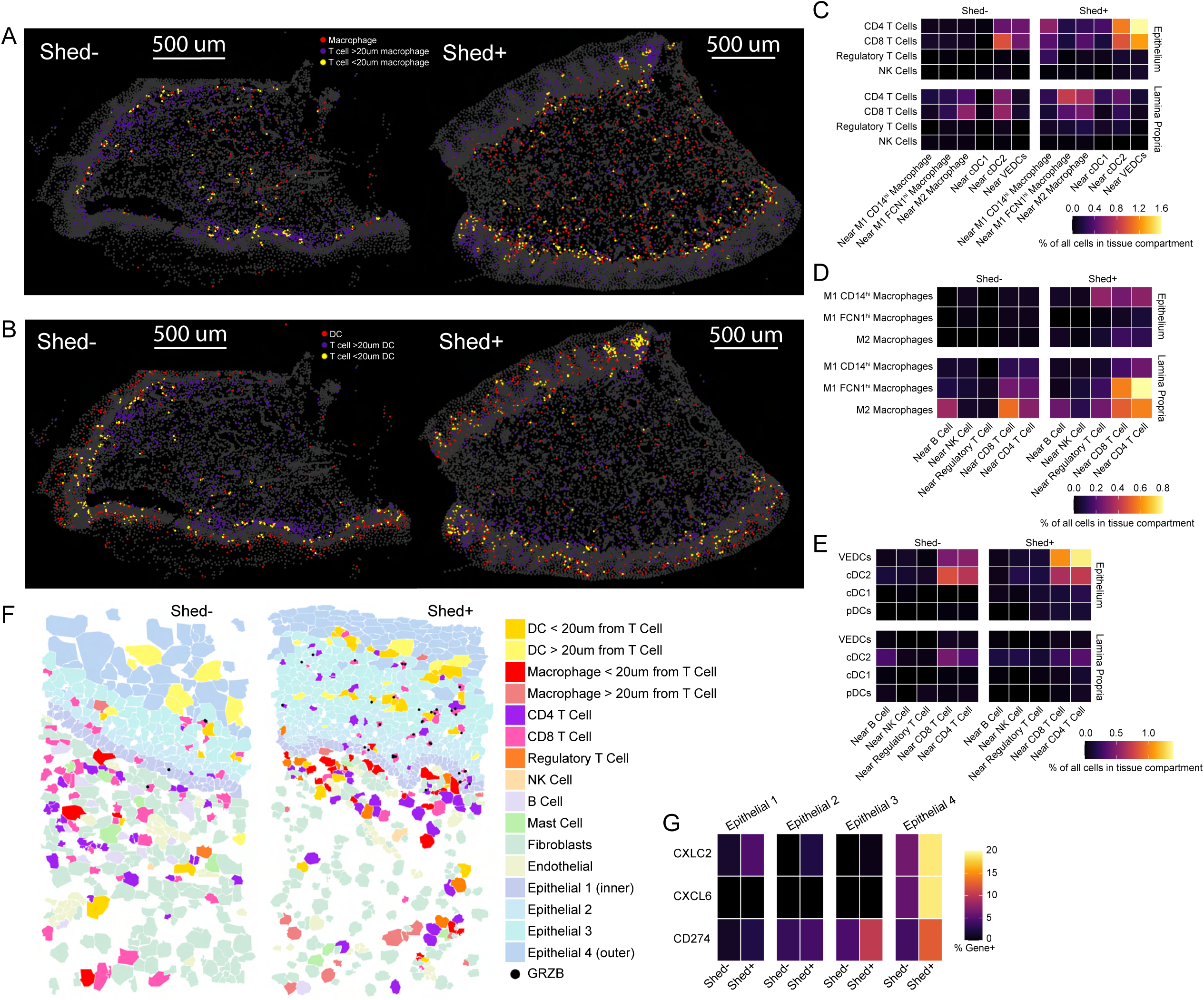
HSV-2 shedding promotes cellular interactions in vaginal epithelium. (A) Representative image of a Shed- and Shed+ VT tissue section highlighting T cells further than 20 μm away from macrophages (purple), macrophages (red) and T cells near macrophages (yellow). (B) Representative image of a Shed- and Shed+ VT tissue section highlighting T cells further than 20 μm away from dendritic cells (purple), dendritic cells (red) and T cells near dendritic cells (yellow). (C) Heat map showing the proportion of T cells within 20 μm of select innate immune cells. Heat map showing the proportion of macrophages (D) and dendritic cell types (E) within 20 μm of T cells stratified by subsets and NK cells. Representative portion of a tissue section from a Shed-(F) and a Shed+ sample to demonstrate cellular interactions that are more frequently observed in Shed+ samples. (H) Heatmap showing CXCL2, CXCL6, and CD274 gene expression by epithelial subtype in Shed- and Shed+ samples.

These findings were confirmed by looking at which cells the macrophages were interacting with, with mostly M2 macrophages interacting with CD8 T cells in the lamina propria in Shed-samples (Figure 7D, Supplemental Figure 7, H and I). In Shed+ samples, most interactions were between FCN1+ M1 macrophages and CD4 T cells in the lamina propria (Figure 7D). Notably, there were also more interactions between FCN1+ M1 macrophages and CD8+ T cells in Shed+ samples (Figure 7D), despite there being a slight decrease in CD8+ T cells in the lamina propria associated with active HSV-2 shedding (Figure 6E). While there were very few macrophage interactions in the epithelium of Shed-samples, there were interactions between M2 and, particularly, CD14hi M1 macrophages and CD4+, CD8+, and regulatory T cells in the epithelium of Shed+ samples (Figure 7D). This data suggests that during HSV-2 shedding, there are slight increases in T cell interactions, particularly CD4+ T cell interactions with M1 macrophage populations in the lamina propria, and that uniquely in the context of active viral shedding, macrophage – T cell crosstalk also occurs in the epithelial layer. Given the different proximities of the macrophage populations from the basal epithelial layer (Figure 6C), this data suggests that in Shed+ samples, the interaction between FCN1hi M1 macrophages and CD4 T cells occurs in the lamina propria, closest to the basal epithelium, and CD14hi M1 macrophages interact with T cells in the epithelium, near the basal epithelium.

Finally, we looked at DC and T cell interactions and found that cDC2 was the dominant subtype which interacted with both CD4 and CD8 T cells in Shed-samples (Figure 7E, Supplemental Figure 7, J and K). In samples from individuals with active HSV-2 shedding, however, this shifts to a larger fraction of VEDC interacting with CD4 T and CD8 T cells, with the addition of cDC1s interacting with CD4 T cells in the epithelium (Figure 7E). This suggests that in Shed+ samples, T cells are being drawn towards VEDCs, which reside in the outer layers of the epithelium.

Through visual examination of spatial interactions, we observe that in the Shed-sample, fewer macrophages are interacting with T cells in the lamina propria, with some CD8 T cells in the epithelial layer, but not far enough to interact with the VEDCs in the outer epithelium (Figure 7F). In contrast, in the Shed+ sample, we observe macrophages interacting with CD4, CD8, and regulatory T cells in close proximity to the basal epithelium, with more CD4 and CD8 T cells in the epithelium interacting with DCs, particularly further towards the outer layers (Figure 7F). This supports our interpretation that during active viral shedding, T cells are being drawn from the lamina propria towards the basal epithelium and interacting with macrophages, then into the outer vaginal epithelium, where they are interacting with VEDCs that express genes associated with maturation and activation.

Finally, we investigated the vaginal epithelial cell compartment for differences in gene expression that may be relevant to the immune responses to HSV-2 shedding. An analysis of the epithelial cells found that a greater proportion of the outer epithelial cells expressed CXCL2, CXCL6, and CD274 in the Shed+ samples (Figure 7G, Supplemental Figure 7L). CD274, or PD-L1, is known to be upregulated when stimulated by interferon in response to viral infection (Schonrich and Raftery, 2019), providing support that viral-immune interactions are occurring in the outer epithelium of the VT during active HSV2 shedding. These inflammatory cues induced by epithelial cells in conjunction with immune cells responding to an active viral infection could drive the mobilization and activation of the DC and macrophages, which may, in turn, play a role in the recruitment and activation of T cells. Altogether, our characterization of immune cell organization and gene expression in the vaginal lamina propria and epithelium in HSV-2 seropositive individuals reveals that active viral shedding associates with changes in immunity within the genital mucosa. Notably, we observed changes in macrophage, dendritic cell, and T cell mobilization toward the epithelial layers, and increased expression of genes related to inflammation, cytotoxicity, activation, and intrinsic regulation.

## Discussion

Previously, many studies investigating immune response dynamics against HSV-2 reactivation have understandably focused on genital lesions, where viral infection and reactivation are known to occur, and described cellular alterations throughout healing. While other studies have analyzed immune cells from those with HSV-2 that were collected from the cervicovaginal lumen via cytobrushes, lavage, or other minimally invasive techniques (Posavad et al., 2015; Shannon et al., 2014), we investigated HSV-2-driven immune responses in VT and CX tissue biopsies. Given the rich immune populations that reside in the deeper tissue layers that contribute to mucosal immunity (Lund et al., 2023), we believe that investigating immune cell populations within the tissue is crucial to unlocking novel insights that can lead to therapeutic interventions. To progress our understanding of how HSV-2 seropositivity correlates with mucosal and circulating immune signatures at a population level, we used advances in tissue cryopreservation and flow cytometry to characterize T cells among a cohort of 232 Kenyan women. Matched anogenital swabs at the same time of biopsy collection and subsequent PCR testing allowed us to identify whether a subset of participants were actively shedding HSV-2 when they provided genital mucosal samples. Recent innovations in subcellular-resolution spatial transcriptomics allowed us to not only address how HSV-2 seropositivity broadly impacts mucosal and circulating immunity but also how episodes of viral shedding are associated with acute immune alterations that may reinforce tissue-specific immune responses in a predominantly asymptomatic cohort. While we did not see shifts in major T cell subsets in circulation or tissue compartments (Figure 1), we did observe phenotypic alteration of VT T cells with minimal observed differences in any of the CX T cell subsets analyzed (Figure 2).

For example, we observed greater CD39 expression on CD8+ T cells in HSV-2 seropositive individuals (Figure 2B). CD39 is an enzyme responsible for converting adenosine triphosphate into adenosine diphosphate and cyclic adenosine monophosphate (Timperi and Barnaba, 2021), and is a marker of exhaustion known to correlate with viral load in chronic infections (Gupta et al., 2015). Its expression is associated with metabolic stress and immune dysfunction, indicative by CD39-expressing cells being less responsive to vaccines and prone to apoptosis (Fang et al., 2016). We also observed a modest increase of CD39 on VT Tconv in HSV-2 seropositive individuals (Figure 2C), though the adjusted results were not significant.

These differences may influence intrinsic metabolic regulatory mechanisms that shape tissue-specific immune responses in the VT. In contrast, we observed a reduction in PD-1 expression on VT Tconv in HSV-2 seropositive individuals (Figure 2C). Both PD-1 and CD39 are associated with exhausted phenotypes (Balanca et al., 2021), so opposing expression patterns in association with HSV-2 seropositivity is surprising. However, what may explain this seesaw effect is shifts in metabolic regulation. PD-1 is associated with deleterious mitochondrial function via recruiting phosphatases and disrupting glycolysis (Kumar and Chamoto, 2021). To compensate, cells expressing PD-1 rely more on oxidative phosphorylation (Bengsch et al., 2016), a metabolic process that CD39 can impair.

We hypothesized that HSV-2 infection, indicated by seropositivity, would have the greatest impact on altering genital immune responses. While it was associated with some alterations in VT T cell phenotypes, HSV-2 seropositivity was surprisingly associated with more differences in the circulating immune signature. A key marker defining this immune signature was increased CCR5 expression on Th1, CD8+ T cells, and Tregs. CCR5-expressing T cells have been well documented to play a critical role in tissue-specific responses, as the chemokine receptor that can help draw T cells into barrier tissue sites. While the frequency of CCR5-expressing cells is dramatically lower in the blood than in the tissue (Figures 2A and 3), elevated CCR5 expression on circulating immune cells may allow them to more readily home to sites of viral reactivation. Mouse models have shown that during HSV-2 re-exposure, macrophages produce CCL5, the ligand for CCR5, thereby leading to the recruitment and retention of CCR5-expressing T cells in the mucosa (Iijima and Iwasaki, 2014). While we do not observe an increased frequency of CCR5 expression in the genital tract mucosal tissue, circulating CCR5-expressing memory T cells may support genital skin-specific immune responses, where CCR5-expressing cells have been identified (Zhu et al., 2009).

In addition to the VT immune signature that we identified in HSV-2 seropositive participants, we also observed signs of intrinsic and extrinsic regulation on circulating T cells. Although the frequency of CD39 expression on T cells is overall lower in the blood than in the tissue (Figures 2 and 3), its frequency of expression was significantly increased on circulating Th1 and CD8+ T cells in HSV-2 seropositive individuals, which may indicate metabolic stress within these cells. Increased CCR5 expression and a modest but insignificant increase in CD39 expression on circulating Tregs may promote Treg activation and greater ability to mediate immunoregulation. While we observe a greater frequency of activated, double-positive CD38+HLA-DR+ Th1 cells, we observe a reduction in their cytotoxic capacity via reduced granzyme B expression in HSV-2 seropositive individuals (Figure 3B). Reduced Granzyme B expression, along with the reduction of several circulating proinflammatory cytokines (Supplemental Figure 2A), could indicate a degree of immunosuppression, perhaps a product of greater regulatory signals provided both intrinsically, via CD39, as well as extrinsically via Treg-mediated suppression.

We hypothesized that the acute and transient nature of active HSV-2 shedding from the genital skin may lead to immune activation in the neighboring VT and be responsible for altering the spatial organization of cells. Similar to our flow cytometry analysis, we saw no broad differences in the immune cell proportions between HSV-2 seropositive vs HSV-2 seronegative samples via spatial transcriptomics. However, comparing Shed+ vs Shed- samples revealed the greatest differences among immune cell subsets and gene expression patterns. Notably, T cells had more detectable CD69, an activation marker promoting tissue retention, and CLCA2, which may also aid tissue retention, specifically by elevating E-cadherin binding in the epithelium (Qiang et al., 2018). Additionally, Shed+ samples had a larger Treg population and more CTLA-4 and IL2R expression on Tregs, which supports Treg suppressive function and cell survival in the tissue. The findings suggest that vaginal T cells may adapt to recurrent viral shedding by promoting regulatory phenotypes that promote disease tolerance (Medzhitov et al., 2012; Schneider and Ayres, 2008; Soares et al., 2017) during HSV-2 shedding.

We also investigated antigen presenting cells to define how their transcriptional profile differs between Shed+ vs Shed-samples. In Shed+ individuals, we observed an innate cell phenotype that associates with proinflammatory responses. Of note, we observed more M1 macrophages with a CD14hi phenotype in Shed+. A greater proportion of these macrophages also expressed CXCL9 and CXCL10, which can promote T cell recruitment to the tissue.

Additionally, in Shed+ samples, we observe significantly more proinflammatory DC with a cDC1 phenotype that express CCR7, LAMP3, and CD83. LAMP3 is upregulated in vaginal epithelium during HSV-2 infection (Nazli et al., 2022) which provides further support that these innate immune cells are likely interacting with HSV-2 virus in the vaginal tissue. We also identify a vaginal epithelial DC-like phenotype we describe as VEDCs which resided in the outer epithelial layer. The VEDC population expresses C15orf48, which was upregulated in Shed+ samples. This gene is elevated in response to inflammatory signals and helps to regulate mitochondrial function when oxidative stress occurs, negatively regulating the immune response (Takakura et al., 2024). C15orf48 expression in VEDCs in Shed+ may contribute to the activation and regulation of inflammatory VEDCs. The switch to proinflammatory phenotypes and upregulation of inflammatory genes is consistent with exposure to viral antigen in the VT during active HSV-2 shedding. There was also upregulation of regulatory immune phenotypes and genes associated with immune regulation, suggesting immune regulatory mechanisms were also being activated in this context of repeated viral exposure.

While there were notable differences when comparing the overall tissue, the greatest insight from this analysis came from assessments of the spatial organization within the tissues and co-localization of immune cells. We observed CD4+ and CD8+ T cell recruitment into the epithelial layer in Shed+ samples, where viral exposure is likely to occur due to anatomical proximity to the genital skin. In Shed-samples, the CD4+ and CD8+ T cells resided in the lamina propria and displayed a resting memory phenotype, including the expression of KLRB1 and CD27, which is expressed by T cells that are not active but primed for recall. We hypothesize that these cells are likely antigen-experienced from previous active shedding events and are retained in the tissue as inactive memory when shedding is not active. In contrast, T cells in Shed+ samples appeared to migrate to the epithelium, where they expressed genes associated with cytotoxicity and activation. The migration and phenotype change in Shed+ samples is evidenced by loss of T cells with a resting memory phenotype in the lamina propria, however, a portion of T cells in the epithelium retained the expression of KLRB1, a marker for memory T cells, decreased expression of CD27, and increased expression of granzyme genes. It should also be noted while this shift was evident for CD8 T cells, there was an additional increase in CD4+ T cells, which had low expression of proliferation genes (MKI67), suggesting these cells have been recruited, possibly from the blood where we saw significantly more CCR5 expression on CD4+ T cells in HSV+ blood samples (Figure 3B). Additionally, greater CD4 T cell localization and activation in the vaginal epithelium of those with active HSV-2 viral shedding may contribute to increased HIV target cells and thus increased HIV susceptibility.

Macrophages and DCs were also more activated in the epithelium of Shed+ samples compared to the epithelium of Shed-samples. Shed+ samples had more M1-like macrophages, likely derived from recruited monocytes due to their expression of FCN1 and CD14 (Sanchez Vasquez et al., 2025; Wang et al., 2024). While FCN1 macrophages were located in the lamina propria, close to the basal epithelium, CD14hi macrophages were increased in the epithelium in Shed+ samples, adding to the inflammatory profile within the epithelium. Additionally, the populations that were found furthest into the epithelium and closest to the vaginal lumen, including cDC1s, VEDCs, and CD14hi M1 macrophages, had upregulation of proinflammatory and maturation genes, which may indicate encounters with viral antigen. Of note, VEDCs expressed the co-stimulatory receptor TNFRSF9 (41BB/CD137), which promotes T cell proliferation, survival, and cytokine production. However, they also expressed the immune checkpoints CD274 (PDL1) (Bryant-Hudson and Carr, 2012) and HAVCR2 (TIM3) (Dixon et al., 2021; Wolf et al., 2020), which negatively regulates T cells, perhaps helping to prevent excessive inflammation.

After discovering various proinflammatory immune cells, both innate and adaptive, we sought to identify cell-cell interactions. We used a 20μm cutoff, previously shown to be robust in identifying cells in direct contact, and used this metric to infer cell-to-cell interactions (Alon et al., 2021). We found T cells in Shed+ were much more likely to be found in proximity to macrophages and DCs (Figure 7C). The recruitment of T cells towards the epithelium may also be augmented by macrophage chemokine production, with the greatest concentration near the basal epithelial layer (Supplemental Figure 6E and F), where most macrophages were located and most T-cell macrophage interactions occurred in Shed+ samples.

HSV-2 shedding was not only associated with the trafficking of immune cells to the epithelium, but also the production of transcripts associated with immunoregulation that may promote disease tolerance and reduce clinical symptoms. Of key importance to regulating inflammation, we observed that CD274 transcripts increased in the outer epithelial cells, and in the DCs and macrophages within the epithelium, where we postulate virus-immune cell interactions and proinflammatory immune responses are most likely occurring in Shed+ samples. CD274, the gene for PD-L1, is upregulated in response to interferon and acts as an extrinsic and intrinsic immune regulator (Hudson et al., 2020). While we observe an increase in epithelial cytotoxic T cells, we simultaneously observe increased regulatory T cell activation (Figure 6H). Additionally, CTLA4+ expression was elevated on T cells within the epithelium of Shed+ samples, suggesting a regulatory response following immune activation. These findings reveal that while proinflammatory immune response may be vital to limiting prolonged viral activation, immunoregulatory networks are likely crucial to balance the tissue response and potentially limit disease pathology, thereby promoting tissue homeostasis. Overall, the shifts that we identified in T cell phenotypes associated with HSV-2 seropositivity and active shedding appear to reflect a balance between inflammation and immune regulation. These phenotypes appear to be amplified during episodes of acute mucosal immune activation associated with HSV-2 reactivation and viral shedding, concurrent with immunoregulatory signals. This regulatory control is likely to be critical during episodic infection, given the known ‘sensing and alarm function’ of T_RM_ (Rosato et al., 2023). As T cells migrate into the epithelial layers of the vaginal mucosa in response to local viral shedding, engagement with their cognate antigen may initiate a robust immune cascade at the tissue site, including bystander activation. In such a setting, mechanisms of immune regulation are essential to preserve tissue integrity and prevent immunopathology.

Our study has several limitations. First, our analysis of T cell populations via flow cytometry had limited power to evaluate phenotypic differences in the rarer cell populations due to a limited number of cells isolated from each mucosal biopsy. Additionally, albeit large, our phenotyping antibody panel was limited, leaving many markers unevaluated. Our analysis of tissue sections via immunofluorescent staining and cellular imaging was limited to only three markers and a limited number of samples that were of sufficient quality for analysis. While our spatial transcriptomics allowed for subcellular resolution, it is a probe-based assay that only evaluated 377 individual mRNA transcripts, with only 121 immune-related genes to define cells and phenotypes, leaving many key genes unchecked. While Xenium does provide good sensitivity, it cannot give true gene counts due to the probe-based design, and all samples provided for analysis were small, which introduces potential bias. Lastly, our clinical data was limited to two clinical visits. PCR testing for HSV-2 shedding was only done on a subset of HSV-2 seropositive participants at one visit, leaving the frequency and history of HSV-2 shedding in the cohort unknown.

In sum, our spatial transcriptomics analysis revealed that the localization, gene expression, and interactions of immune cells are profoundly altered by HSV-2 viral shedding. The vaginal tissue response to viral shedding may be dependent not only on T cell recruitment and cytotoxic activation, but also regulatory mechanisms including expression of intrinsic immune checkpoint factors to balance a memory tissue response that can quickly contain viral shedding episodes while also reducing the occurrence of symptomatic herpes infection and adverse health outcomes associated with HSV-2 infection.

## Materials and Methods

### Participants, samples, and data collection

Samples for this analysis were provided by a subset of participants from the Kinga Study (Clinicaltrials.gov ID# NCT03701802). In total, the Kinga Study enrolled 406 heterosexual Kenyan couples to study how sexually transmitted infections and other exposures affect genital tract and systemic immune responses. Of the 406 couples enrolled, this analysis includes N=135 HSV-2 seronegative and N=97 HSV-2 seropositive individuals identified as female at birth (Table 1). Inclusion criteria for this analysis included those with a conclusive (positive or negative) HSV-2 serologic test result and a negative HIV-1/2 serologic test. Positive calls for the HerpeSelect-2 EIA (Focus Technologies, Cypress CA) assay used an index value cut-off of 3.4 (and a negative value below 1.8) to improve test specificity (Ashley-Morrow et al., 2004; Gamiel et al., 2008; Golden et al., 2005; Laeyendecker et al., 2004). Additionally, only samples that were provided by participants at their enrollment visit were analyzed here.

Eight sample types were requested (in addition to diagnostic clinical samples) from each Kinga Study participant for immune characterization studies. These include two 3-mm VT tissue biopsies and two 3-mm CX tissue biopsies. Each biopsy was cryopreserved with one VT and one CX biopsy cryopreserved in optimal cutting temperature (OCT) and used for cryosectioning and immunofluorescent imaging and/or spatial transcriptomics analysis, and one VT and one CX biopsy cryopreserved for high-parameter flow cytometry analysis. Fractionated PBMC samples were also collected and cryopreserved for high-parameter flow cytometry analysis. CVT fluid was collected by menstrual cup (Softcup^®^) and, along with serum samples, analyzed for the presence and concentration of soluble immune factors. Additionally, an anogenital swab was collected at the study visit for PCR testing to detect active HSV-2 viral shedding. Using a Dacron swab, the anogenital region was sampled, which includes the cervix and vagina, labia majora and minora and perineum/perianal region. Sample collection and clinical testing procedures for the Kinga Study, including HIV-1/2 and HSV-2 serology testing methods, were performed in the same manner as previously described (Gamiel et al., 2008; Golden et al., 2005; Laeyendecker et al., 2004; MacLean et al., 2025).

### Immunofluorescent microscopy analysis

A subset of participants in this analysis provided a VT or CX sample that was sectioned and analyzed for the quantity and spatial arrangement of CD3+, CD4+, and CCR5+ cells. This subset includes only participants who were not exposed to HIV and had no BV (a Nugent score of 0-3). Biopsies were collected by clinicians at the study site using Tischler forceps at the lateral VT wall and/or the ectocervical os, embedded in OCT compound, and cryopreserved on dry ice. Laboratory analysis, including tissue processing, staining, imaging, and other assay components, including tissue section image analysis, was done exactly as described previously (MacLean et al., 2025). The codes and an extended description of the GUI used to analyze immunofluorescent images in MATLAB (R2022b) can be found at github.com/FredHutch/Kinga_Study_BV_MacLean (MacLean et al., 2025).

### High-parameter flow cytometry analysis

Participants also provided VT and CX biopsies cryopreserved for high-parameter flow cytometry analysis. Study site clinicians collected these biopsies using Tischler biopsies at the VT wall or cervical os. Biopsies were placed in cryovial with 4% fetal bovine serum at 4°C, transported briefly to the lab, then cryopreserved overnight at -80°C in dimethyl sulfoxide (DMSO), and transferred to liquid nitrogen for long-term storage as described previously (Hughes et al., 2018). PBMCs were also collected, processed, and cryopreserved in DMSO. All processing for genital tissues and PBMC samples was done previously as described in detail (MacLean et al., 2025). Samples were then shipped to the University of Washington for further analysis. Cryopreserved tissue biopsies and PBMC samples were quickly thawed, processed, and stained with fluorescently labeled antibodies as previously described (MacLean et al., 2025).

After isolation, cells were incubated with UV Blue Live/Dead reagent for 30 minutes at room temperature and then stained with the antibodies and protocol previously described (MacLean et al., 2025). Analysis was performed using Flowjo software. Gating analysis was also replicated as described by Maclean et al. 2025, with a 25 cell minimum required to proceed to any downstream gate. The proportion of each cell type was then calculated by dividing the count of a phenotype by the count of its parent gate.

### Cytokine and chemokine analysis

A menstrual cup inserted into the vaginal canal of participants who were not actively menstruating was used to collect CVT fluid for analysis of cervicovaginal soluble immune factors. Serum samples were also collected to analyze circulating soluble immune factors. CVT fluid and serum samples were processed, cryopreserved, and shipped to the University of Washington and then to Eve Technologies (Calgary, Alberta, Canada). The detection and quantification of chemokines and cytokines were done using the Human Cytokine Array/Chemokine Array 71-403 Plex Panel (Eve Technologies, HD71). All collection and processing were done as described previously (MacLean et al., 2025).

### Spatial Transcriptomics Sampling and Processing

In addition to previously described clinical testing, a subset of HSV-2 seropositive individuals provided an anogenital swab at the same time as biopsy collection for the detection of viral DNA via PCR testing previously described by Jerome *et al*. (Jerome et al., 2002). PCR was performed on a total of 85 anogenital swabs at the University of Washington Virology Lab. N=12 swabs were positive for HSV-2, indicating that these participants were actively shedding HSV-2 virus at the same time as biopsy collection. A subset of 3 individuals with a positive HSV-2 PCR result, 2 individuals HSV-2 seropositive but with a negative HSV-2 PCR result, and 6 HSV-2 seronegative participants provided fresh VT biopsies, which were cryopreserved in OCT and included in the spatial transcriptomics analysis. Of the five HSV-2 seropositive participants included here, only 1 reported having sores in the last three months. All 5 HSV-2 seropositive participants showed no signs of active genital herpes during their clinical exam at the visit.

Fresh frozen VT biopsies from this subset of participants were prepared according to the 10x Genomics tissue preparation protocol (CG000579, Rev C). OCT blocks were cryosectioned at 10μm, adhered to equilibrated Xenium slides, and stored at -80°C for later processing. Tissue sections were then fixed in paraformaldehyde and permeabilized following the protocol CG000581 Rev C. Probe hybridization of 377 genes in the Xenium Human Multi-Tissue and Cancer panel (10x Genomics, 2023), ligation, and amplification were performed as instructed in CG000582 Rev E, followed by nuclei staining and removal of background fluorescence. Imaging and decoding of fluorescent probe optical signatures were performed onboard the 10x Genomics Xenium platform (software version 1.7.6.0 and analysis version xenium-1.7.1.0) with additional analysis to detect the abundance and localization of mRNA transcripts.

### Xenium Analysis

#### Data preprocessing and cell segmentation

Cell segmentation was initially performed with 10x Xenium onboard cell segmentation (10x Genomics). DAPI-stained nuclei from the DAPI morphology image were segmented, and boundaries were consolidated to form nonoverlapping objects. Next, we used Proseg v1.1.8 (Jones et al., 2025) cell segmentation algorithm, which uses cell morphologies identified as described above, plus the spatial distribution of transcripts to determine cell boundaries. Samples were then processed with Xenium Ranger v 3.0.1.

#### Quality filtering and data preprocessing

Data was loaded and analyzed using R version 4.3.1. For each sample, Xenium generated an output file of transcript information, including x and y coordinates, corresponding gene, assigned cell and/or nucleus, and quality score. Low-quality transcripts (quality value (QV) < 20) and transcripts corresponding to blank probes were removed. Seurat v5 (Hao et al., 2024) was used to perform further quality filtering and visualization on gene expression data. A Seurat object was created for each sample, containing probe counts, metadata, centroids, segmentations, and molecules. Cells that contained ≥10 transcripts corresponding to ≥5 unique genes were retained. Individual samples were normalized individually, and a single merged Seurat object containing a total of 142,903 cells was created for all samples for scaling and PCA. To account for interpatient sample-specific variability, samples were integrated using Seurat RPCA integration. The integrated object was used to generate UMAP and clustering using Seurat FindClusters with a resolution of 0.5.

#### Cell type identification

To annotate cell clusters, SingleR v2.4.1 (Aran et al., 2019)was employed to compare against 6 different references encompassing both tissue and immune cell atlases available through the SingleR celldex v1.12.0 package (Human Primary Cell Atlas, Database of Immune Cell Expression, ENCODE Blueprint data, ImmGen 28 cell, Monaco Immune data, Mouse Cell Atlas (converted to human genes)). 12 main clusters were identified and annotated. The T cell and macrophage/DC clusters were subsetted and regressed for genes associated with their spatial proximity, such as epithelial and fibroblast genes, and sub-clustered to identify T cell and myeloid subsets.

#### Gene expression

For analysis, HSV+ samples were defined as Shed+ or Shed- as described above. Seurat’s FindMarkers function was used to find differentially expressed genes for each subtype between Shed+ and Shed- samples (LFC>1 and padj<0.05). Due to the probe-count-based approach of the Xenium platform, which does not reflect true gene counts, we considered a cell positive for a gene if it contained 1 or more normalized counts, and calculated the percentage of cells positive for a gene as a comparator. Heatmaps were generated using ggplot2 v3.5.1(Wickham, 2016).

#### Identification of tissue compartments

For spatial analyses, individual samples were resubsetted from the merged Seurat object. To discriminate the localization of immune cells between the epithelium and lamina propria, Seurat’s BuildNicheAssay was used on each sample with niches, k = 2, and neighbors, k = 5, to define 2 niches with enough definition to also identify the pockets of lamina propria which bud into the epithelial layer. Cells were designated to either niche, and subsequent analysis compared Shed+ and Shed- samples by niche. To calculate the proportion of a specific cell type, the number of cells was divided by the total number of cells in each niche, per sample. This was done to account for sampling differences of the tissue sections, which had differing numbers of cells and skewing due to the amount of epithelial vs lamina propria tissue captured.

#### Cell distances

To calculate the distance of cells from the border between the two niches, cell coordinates were extracted for each sample, then for each cell, the nearest inner epithelial cell was identified using the RANN package v2.6.1 (Mount, 2024), the distance was calculated, and each cell was assigned a distance value (μm).

#### Density

To calculate density, cell centroids were extracted for each sample and placed into 40 μm bins based on their coordinates. The number of cells of interest was counted for each bin, and bin indices were converted to spatial positions. The density was calculated as the number of cells per bin/area of the bin, and averaged to give cells per um^2^. Due to the small sample size, statistical significance between groups was assessed using a non-parametric, two-sided permutation test (n = 10,000 permutations), comparing the difference in means.

#### Co-localization

To determine which cells were in direct contact, a cut-off of 20 μm was used, and cells within that radius of a cell of interest were determined to be a “near” cell. A 20 μm cut-off was selected as lymphocytes are typically 10 μm in diameter, and macrophages are 20 μm in diameter, and the distance used is between cell centroids (the center of the cell). We therefore reasoned and confirmed that a distance of 20 μm between centroids would capture direct neighboring cells only. A previous study also found that 20 μm was robust in identifying cell-cell interactions (Alon et al., 2021). Cell centroids were extracted and nearest neighbors calculated using frNN from the dbscan package v 1.1-12 (Hahsler et al., 2019), with a neighborhood radius of 20. For each cell type of interest, the cell indices of interest were extracted along with the indices of the nearby cells. Cells that were identified as nearby cells were added to the metadata as “near” the cell of interest. To calculate the frequency of cell interactions in each niche, the number of specific cell interactions, classified as *subset of interest* *near_cell of interest* was divided by the total cells in each niche, per sample.

### Statistics

We compared CD3+, CD3+CD4+, and CD3+CD4+CCR5+ cell densities from immunofluorescent imaging experiments between HSV-2 seropositive vs seronegative participants using the Wilcoxon rank sum test. For flow cytometry experiments, we used rank-based regression, a nonparametric method robust to outliers (McKean and Hettmansperger, 1978), to compare the percentage of specific T cell phenotypes. For soluble immune factor analysis, we first determined the proportion of total samples that were at least 80% quantifiable for each cytokine/chemokine measured. For those cytokines/chemokines with at least 80% quantifiable detection, we imputed out-of-range values by randomly selecting a value between the lowest observed value and half the lowest observed value (if out-of-range low), or by selecting the largest observed value (if out-of-range high). We then estimated differences in mean log cytokine/chemokine concentrations from both serum and CVT fluid using linear regression. If fewer than 80% of samples were quantifiable, then we categorized each value as either detected or undetected. Next, we estimated the odds ratio for the effect of HSV-2 on the detection of those given cytokines/chemokines that were <80% detectable using logistic regression.

For flow cytometry and soluble immune factor analysis of genital tract samples (VT and CX biopsy and CVT fluid), results were adjusted for hormonal contraceptive use (yes, no and menstruating, no and amenorrheal, or unknown for a small number of samples), bacterial vaginosis (Nugent score: 0-3 negative, 4-6 intermediate, 7-10 positive, or unknown for a small number of samples)(Nugent et al., 1991), HIV exposure (HIV status of sexual partner), number of unprotected sex acts in the last thirty days (continuous), and age (continuous variable). Results from PBMC and serum samples were adjusted for hormonal contraceptive use and age only.

This is an exploratory analysis; we did not adjust our results for multiple comparisons. We considered all nominal p-values less than or equal to 0.05 as significant. Statistical analysis was done using R version 4.3.3.

### Study Approval

All participants provided written informed consent using documents reviewed and approved by the University of Washington institutional review board, and the Scientific and Ethics Review Unit of the Kenya Medical Research Institute.

### Data availability

All supporting data are included in the supplemental figures, tables, and/or are available upon reasonable request to the senior authors. The spatial transcriptomics data generated during this study will be deposited in GEO. The custom R code for spatial analyses will be available on GitHub.

## Supporting information

Supplemental Acknowledgments

## Author contributions

FM, JBG, JLS, SCV, NP, ICT, LW, and DS conducted the experiments. FM, RZ, ATT, JLS, AS, and KKT analyzed data. MM provided reagents. RZ, JD, LKS, and KRJ contributed analysis methods. BHC, KN, NRM, JRL, and JML designed the research study. JRL, KKT, AE, EN, JLM, and NRM provided supervision. FM, RZ, JTS, RSM, EN, JRL, and JML wrote the first draft of the manuscript. All authors edited and approved the manuscript.

## Acknowledgments

We thank all study volunteers for their participation in the Kinga Study and their willingness to provide many different samples. Additionally, we thank the members of the Kinga Study team and the Lund and Prlic labs for their helpful discussions on experimental findings and manuscript preparation throughout this process. Additionally, we thank the Fred Hutchinson Cancer Center Cellular Imaging Shared Resource, supported by the Fred Hutchinson Cancer Center Cellular Imaging Core Facility (RRID:SCR_022609) of the Fred Hutch/University of Washington/Seattle Children’s Cancer Consortium (P30 CA015704), for assistance with microscopy and image analysis. SV was supported by T32 AI007509, ICT was supported by T32 AI007140, and LW was supported by T32 AI083203. See Supplemental Acknowledgments for Kinga Study consortium details. This work was supported by the following grants from the National Institutes of Health: R01 AI131914 (to J.M.L and J.R.L.), R01 AI141435 (to J.M.L.), R01 HD114505 (to R.S.M. and J.M.L.), and R01 AI129715 (to J.R.L).

## Figure Legends

**Supplemental Figure 1 - Circulating T cell phenotypes associated with HSV-2 seropositivity** Heatmaps showing the frequency of CD8+ T cell, Tconv, Treg, Th1, and Th17 phenotypes in PBMC samples from HSV-2 seropositive and seronegative individuals. Significant results comparing HSV-2 seropositive vs seronegative, using an adjusted rank regression model that adjusted for hormonal contraceptive use and age, are boxed in blue. N=130 HSV-2 seronegative and N=92 HSV-2 seropositive.

**Supplemental Figure 2 – Soluble immune factor analysis on serum from HSV-2 seropositive and seronegative individuals**

Soluble immune factors were measured from N = 97 HSV-2 seropositive and N = 134 HSV-2 seronegative serum samples. (A) The adjusted 95% confidence is shown. Results that are significant (adjusted P < 0.05) are in red. (B) For soluble immune factors were fewer than 80% of samples were detectable, a dichotomous outcome (present vs not present) was evaluated. The adjusted 95% confidence for the odds ratio analysis is shown. Serum analyses are adjusted for hormonal contraceptive use and age.

**Supplemental Figure 3 – Soluble immune factor analysis on CVT fluid from HSV-2 seropositive and seronegative individuals**

Soluble immune factors were measured from N = 82 HSV-2 seropositive and N = 114 HSV-2 seronegative cervicovaginal fluid samples. (A) The adjusted 95% confidence is shown. Results that are significant (adjusted P < 0.05) are in red. (B) For soluble immune factors were fewer than 80% of samples were detectable, a dichotomous outcome (present vs not present) was evaluated. The adjusted 95% confidence for the odds ratio analysis is shown. Cervicovaginal fluid samples comparisons are adjusted for hormonal contraceptive use, age, semen exposure, HIV exposure, and bacterial vaginosis via Nugent score.

**Supplemental Figure 4 – Spatial identity of vaginal immune cells based on HSV-2 shedding and HSV-2 serostatus**

11 samples were included in the spatial transcriptomics analysis. This includes 6 HSV-2 seronegative, 2 HSV-2 seropositive, anogenital HSV-2 PCR negative, and 3 HSV-2 seropositive, anogenital HSV-2 PCR positive (A). (B) The number of cells captured per sample and the proportion of each cell subset per sample is depicted graphically. The biopsy size and number of cells acquired from each sample was variable, but each cell population was represented in each tissue, and the proportion of the cell types was relatively stable.

**Supplemental Figure 5 – Gene expression in immune cell subsets from HSV-2 Shed- and Shed+ individuals**

Violin plots showing the scaled expression of genes identified as differentially expressed between Shed+ and Shed- in (A) T cell subsets and NK cells, (B) macrophage subsets, and (C) dendritic cell subsets.

**Supplemental Figure 6 – Tissue location of immune cell subsets varies based on HSV-2 shedding status**

(A) The total T cell density in each tissue compartment, separated by Shed- vs Shed+, calculated via spatial transcriptomics. (B) The CD3+CD4+ density in the epithelium divided by the total CD3+ density in the tissue, as measured by immunofluorescent staining and cellular imaging, to account for differences in total cellularity and tissue size across samples. Each dot represents an individual sample, the bar represents the median. Darker shaded dots correspond to the sections from the same samples as those used in the Xenium experiment. N Shed- = 6, N Shed+ = 5.

Comparison using Mann-Whitney test, P value displayed. (C) Bar plot of total gene expression to genes shown in the heatmap of Figure 6F. (D) The total macrophage and subset density in each tissue compartment, comparing Shed- vs Shed+, calculated via spatial transcriptomics. (E) A correlation graph between macrophage CXCL10 expression and distance to the nearest inner/basal epithelial cell stratified by Shed- vs Shed+. (F) A correlation graph between macrophage CXCL9 expression and distance to the nearest inner/basal epithelial cell, stratified by Shed- vs Shed+. (G) The total dendritic cell and subset density in each tissue compartment, comparing Shed- vs Shed+, calculated via spatial transcriptomics. Comparisons for density made using a non-parametric permutation test and for correlation graphs using Pearson correlation coefficients within each niche and Shed group. Statistical significance of each chemokine correlation was assessed using Pearson’s correlation test and corrected for multiple comparisons using FDR method.

**Supplemental Figure 7 – Immune cell organization in the vaginal epithelium and lamina propria in presence or absence of HSV-2 shedding episode**

Representative image of (A) T cells (pink), (B) macrophages (red), or (C) dendritic cells (yellow), and any cell that colocalized near them (blue). (D) The proportion of T and NK cells near macrophages in either tissue compartment in Shed- and Shed+. (E) The density of T cells that colocalized with macrophages in Shed- vs Shed+. Each dot represents an individual sample. Comparison made using a non-parametric permutation test, P value displayed. (F) The proportion of T and NK cells near dendritic cells in either tissue compartment in Shed- and Shed+. (G) The density of T cells that colocalized with dendritic cells in Shed- vs Shed+. Each dot represents an individual sample. Comparison made using a non-parametric permutation test, P value displayed. (H) Bar plot showing the percentage of macrophages near T cell subsets in Shed- vs Shed+ by tissue compartment as a percentage of total macrophages in the tissue compartment. (I) Bar plot showing the percentage of macrophage subsets near a T cell in Shed- vs Shed+ by tissue compartment as a percentage of total cells in the tissue compartment. (J) Bar plot showing the percentage of dendritic cells near T cell subsets in Shed- vs Shed+ by tissue compartment as a percentage of total macrophages in the tissue compartment. (K) Bar plot showing the percentage of dendritic cell subsets near a T cell in Shed- vs Shed+ by tissue compartment as a percentage of total cells in the tissue compartment. (L) Violin plots showing the scaled expression of genes identified as differentially expressed between Shed+ and Shed- in epithelial cell subsets.

