## Supplemental Figure 1 for "Genital herpes shedding episodes associate with alterations in the spatial organization and activation of mucosal immune cells"

**PBMC CD8+ T Cell Phenotypes**

| | CCR7+CD45RA+ | CCR7+CD45RA- | CCR7-CD45RA+ | CCR7-CD45RA- | CCR5+ | CCR6+ | CD161+ | CD38+ | CD39+ | CD69+CD103+ | CD101+ | CXCR3+ | Granzyme B+ | HLA-DR+ | CD38+HLA-DR+ | PD-1+ | TCF-1+ | PD-1+TCF-1- | T-bet+ | CD161+CCR6+ |
|---|---|---|---|---|---|---|---|---|---|---|---|---|---|---|---|---|---|---|---|---|
| HSV-2 Seronegative | **41.1** | 4.3 | **31.2** | 18.2 | **20.7** | 2.4 | 13.6 | 8.8 | **1.2** | 0.1 | 27.3 | 47.7 | 33.7 | 16.4 | 2.7 | 7.0 | 56.9 | 4.0 | 37.7 | 1.3 |
| HSV-2 Seropositive | **37.8** | 3.7 | **35.8** | 19.6 | **26.6** | 2.6 | 11.5 | 7.3 | **1.5** | 0.1 | 31.8 | 53.4 | 34.7 | 20.5 | 3.1 | 7.5 | 54.8 | 4.1 | 38.2 | 1.1 |

**PBMC Tconv Phenotypes**

| | CCR7+CD45RA+ | CCR7+CD45RA- | CCR7-CD45RA+ | CCR7-CD45RA- | CCR5+ | CCR6+ | CD38+ | CD39+ | CD69+ | CD69+CD103+ | CD101+ | CTLA4+ | CXCR3+ | Granzyme B+ | HLA-DR+ | CD38+HLA-DR+ | PD-1+ | TCF-1+ | PD-1+TCF-1- | Th1 | Th17 |
|---|---|---|---|---|---|---|---|---|---|---|---|---|---|---|---|---|---|---|---|---|---|
| HSV-2 Seronegative | 37.6 | 20.0 | 7.1 | 34.0 | 12.4 | 13.8 | 21.7 | 3.4 | 0.9 | 0.0 | 4.4 | 1.3 | 21.1 | **2.9** | 4.9 | 1.4 | 9.3 | 79.0 | 4.2 | 5.0 | 4.8 |
| HSV-2 Seropositive | 34.8 | 19.4 | 6.6 | 34.8 | 14.0 | 14.4 | 19.0 | 3.8 | 0.9 | 0.0 | 4.9 | 1.4 | 23.9 | **2.3** | 5.1 | 1.5 | 9.6 | 77.4 | 4.0 | 4.8 | 4.8 |

**PBMC Treg Phenotypes**

| | CCR7+CD45RA+ | CCR7+CD45RA- | CCR7-CD45RA+ | CCR7-CD45RA- | CCR5+ | CCR6+ | CD38+ | CD39+ | CD69+ | CD69+CD103+ | CD101+ | CTLA4+ | CXCR3+ | Granzyme B+ | HLA-DR+ | CD38+HLA-DR+ | PD-1+ | TCF-1+ | PD-1+TCF-1- | T-bet+ | CCR6+CD161+ |
|---|---|---|---|---|---|---|---|---|---|---|---|---|---|---|---|---|---|---|---|---|---|
| HSV-2 Seronegative | 12.4 | 9.8 | 14.3 | 61.4 | **26.6** | 12.6 | 16.3 | 41.7 | 1.5 | 0.0 | 11.5 | 19.0 | 18.0 | 0.8 | 26.1 | 4.6 | 8.4 | 34.3 | 6.2 | 1.8 | 2.9 |
| HSV-2 Seropositive | 10.6 | 8.5 | 12.3 | 64.6 | **30.5** | 13.7 | 15.3 | 48.1 | 1.4 | 0.0 | 13.5 | 18.3 | 20.9 | 0.8 | 26.5 | 4.5 | 9.4 | 32.1 | 6.2 | 1.9 | 3.2 |

**PBMC Th1 Phenotypes**

| | CCR7+CD45RA+ | CCR7+CD45RA- | CCR7-CD45RA+ | CCR7-CD45RA- | CCR5+ | CD38+ | CD39+ | CD101+ | CTLA4+ | CXCR3+ | Granzyme B+ | HLA-DR+ | CD38+HLA-DR+ | PD-1+ | TCF-1+ |
|---|---|---|---|---|---|---|---|---|---|---|---|---|---|---|---|
| HSV-2 Seronegative | 5.7 | 7.8 | 8.6 | 72.2 | **66.3** | 12.8 | **3.9** | 14.1 | 2.5 | 30.2 | **57.4** | 23.7 | **6.5** | 40.9 | 32.7 |
| HSV-2 Seropositive | 6.2 | 6.8 | 7.3 | 76.0 | **73.9** | 13.3 | **4.3** | 15.8 | 2.9 | 42.2 | **42.9** | 27.1 | **7.7** | 37.3 | 35.5 |

**PBMC Th17 Phenotypes**

| | CCR7+CD45RA+ | CCR7+CD45RA- | CCR7-CD45RA+ | CCR7-CD45RA- | CCR5+ | CD38+ | CD39+ | CD101+ | CTLA4+ | CXCR3+ | Granzyme B+ | HLA-DR+ | CD38+HLA-DR+ | PD-1+ | TCF-1+ |
|---|---|---|---|---|---|---|---|---|---|---|---|---|---|---|---|
| HSV-2 Seronegative | 2.7 | 33.2 | 0.9 | 61.3 | 30.3 | 13.2 | 7.4 | 5.3 | 4.6 | 35.6 | 0.5 | 6.1 | 2.1 | 15.3 | 75.8 |
| HSV-2 Seropositive | 3.0 | 32.3 | 1.1 | 61.9 | 30.3 | 11.5 | 7.8 | 6.3 | 4.3 | 34.0 | 0.4 | 6.3 | 2.6 | 15.4 | 72.8 |

[box] = $P_{adj} < 0.05$

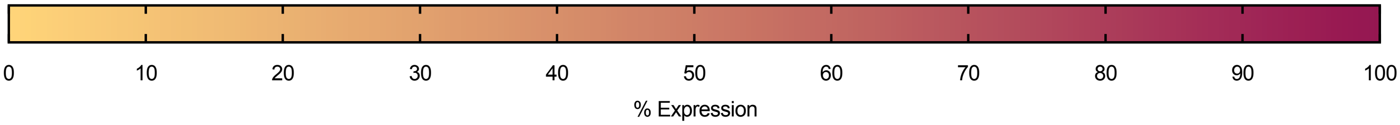
