## Supplementary figures and images for "Genital herpes shedding episodes associate with alterations in the spatial organization and activation of mucosal immune cells"

### Supplemental Figure 2

##### Supplemental Figure 2

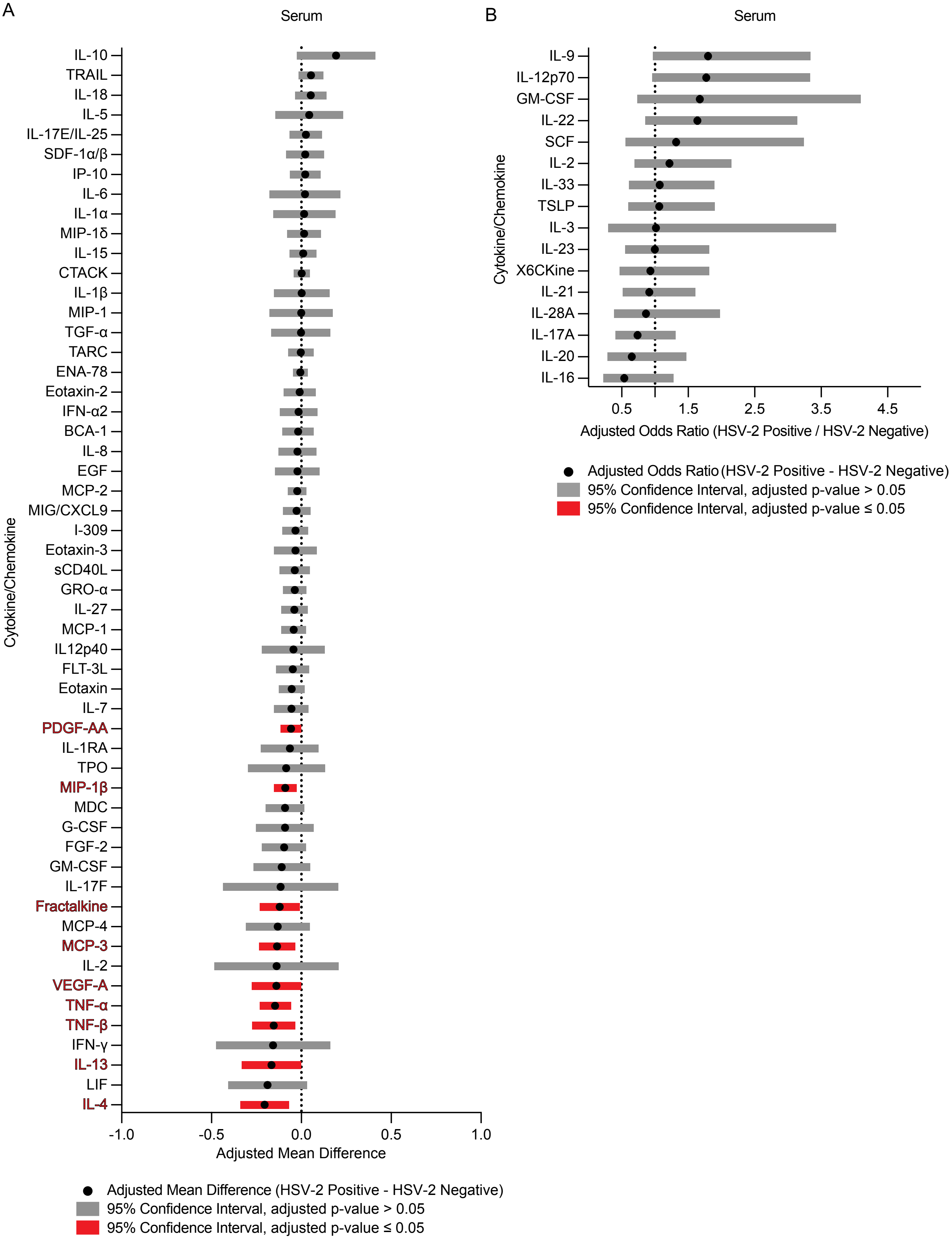

### Supplemental Figure 3

#### Supplemental Figure 3

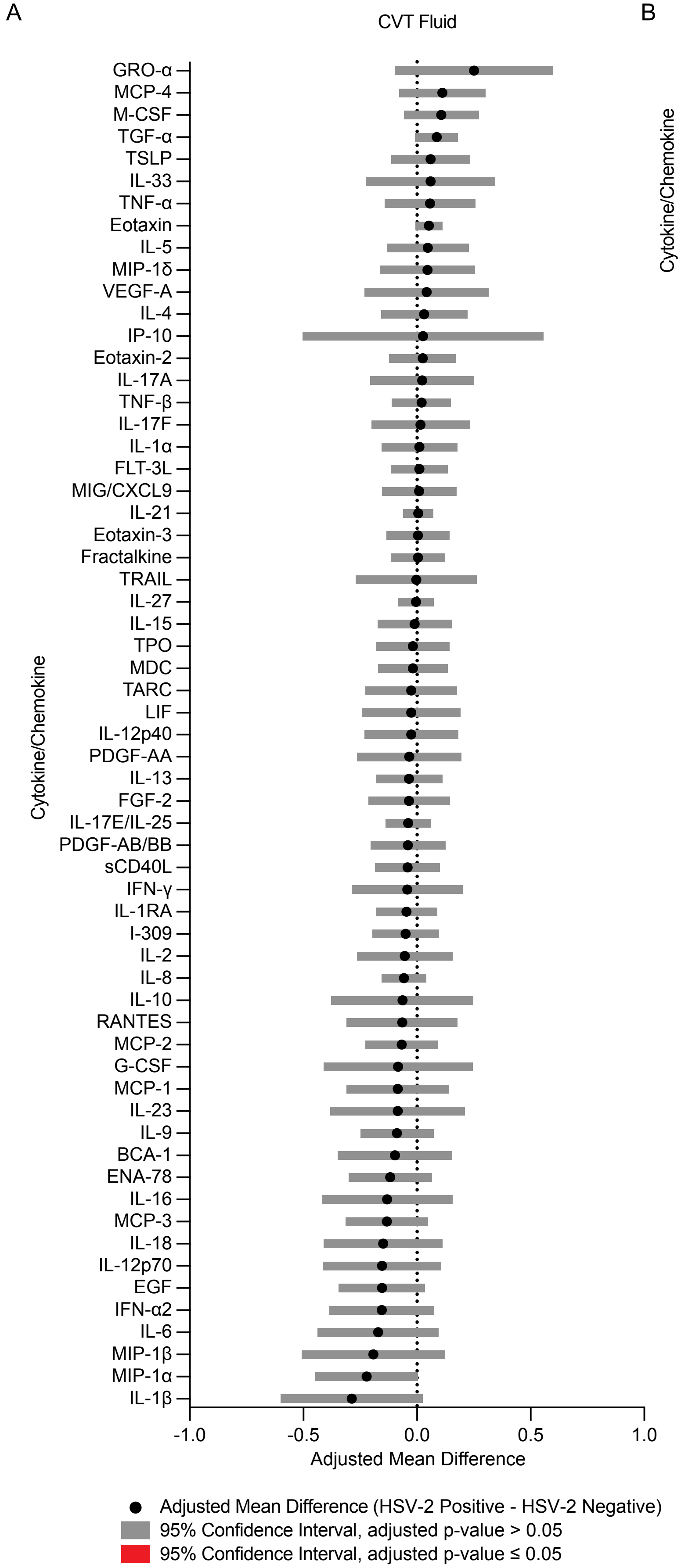


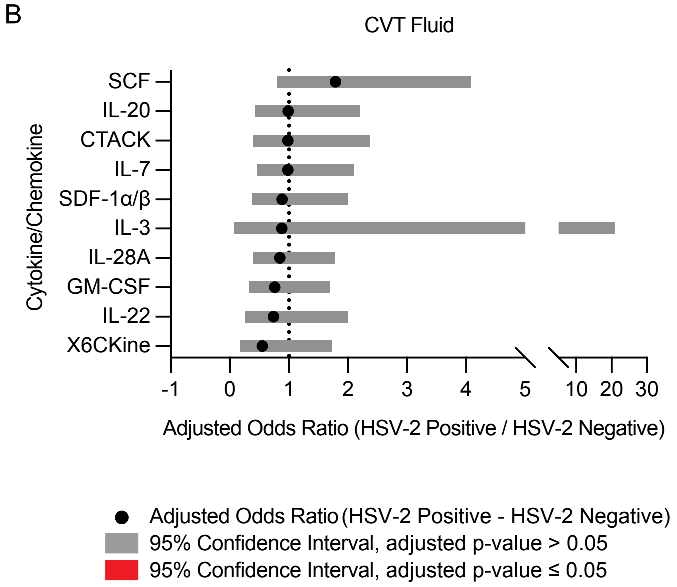

### Supplemental Figure 4

Supplemental Figure 4

A

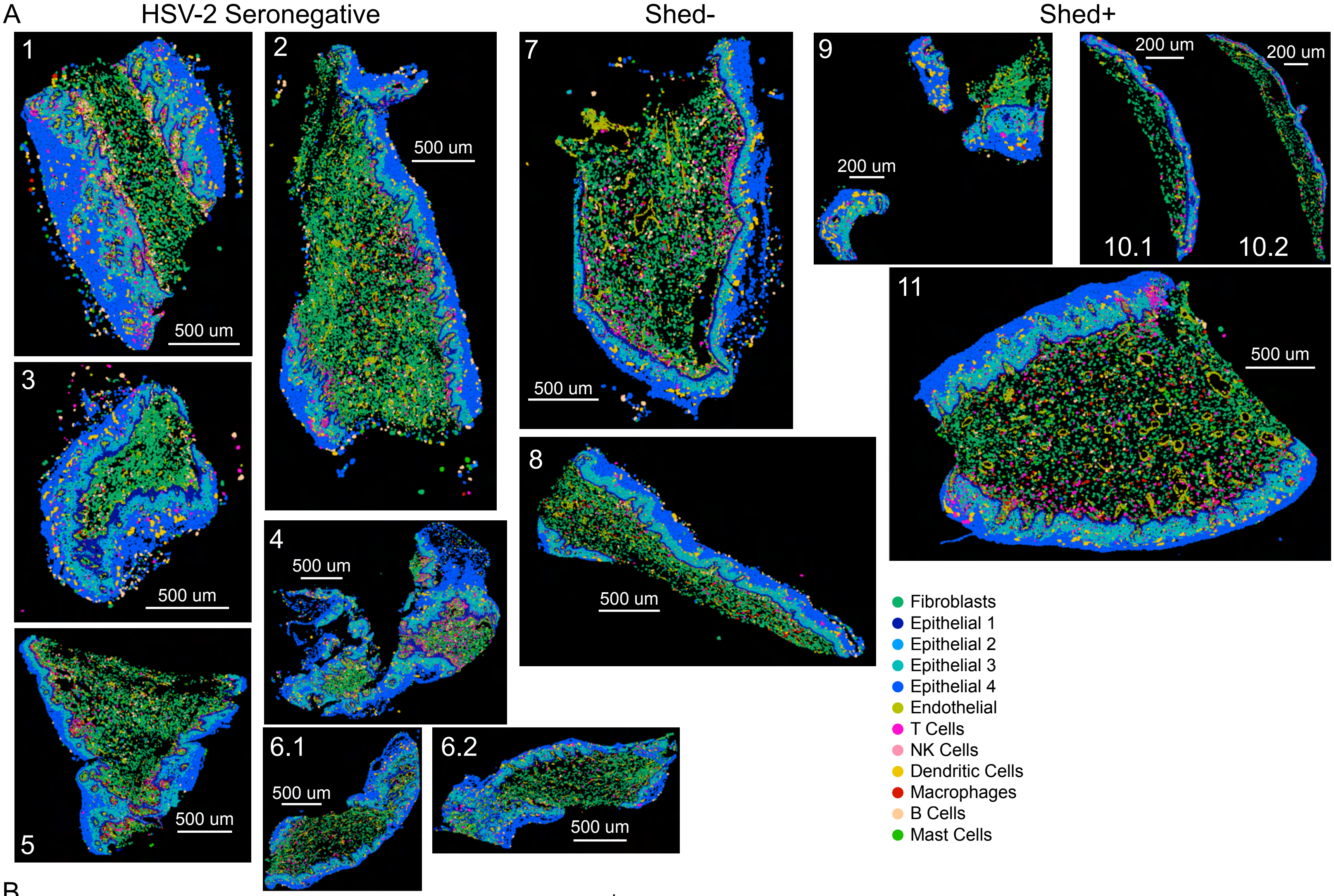


B

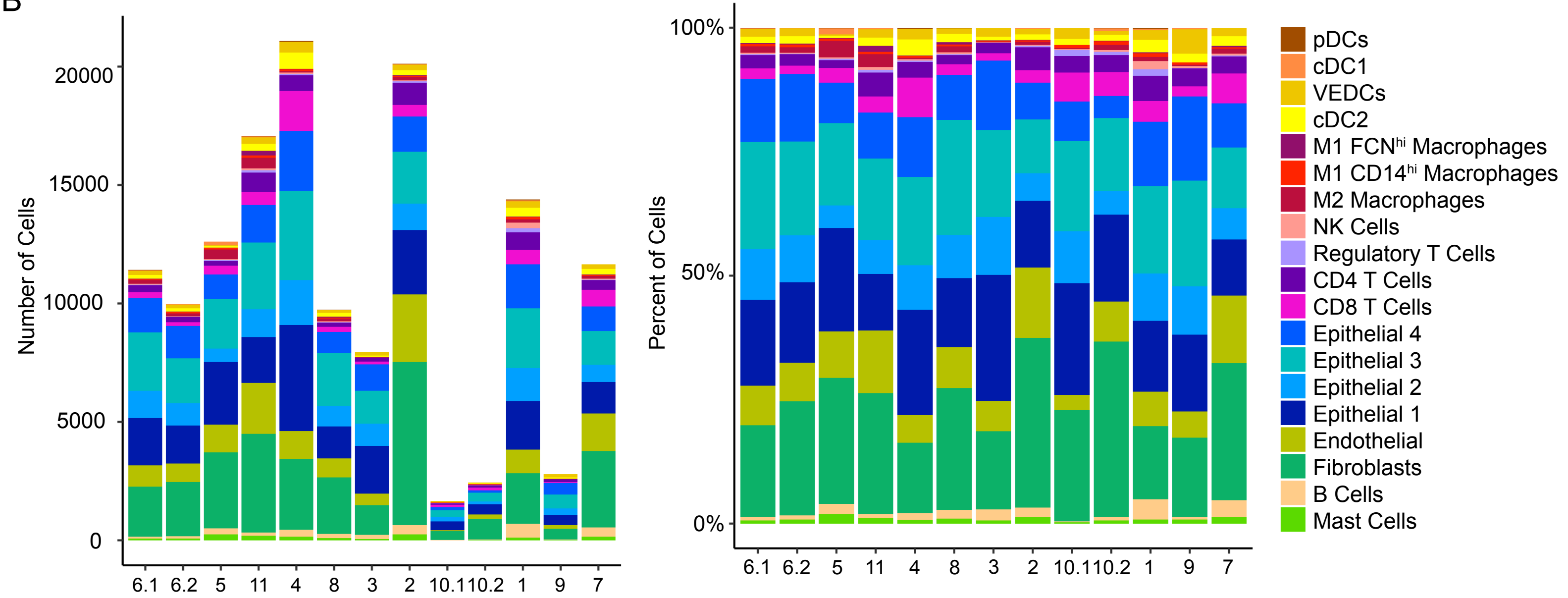

### Supplemental Figure 5

Supplemental Figure 5

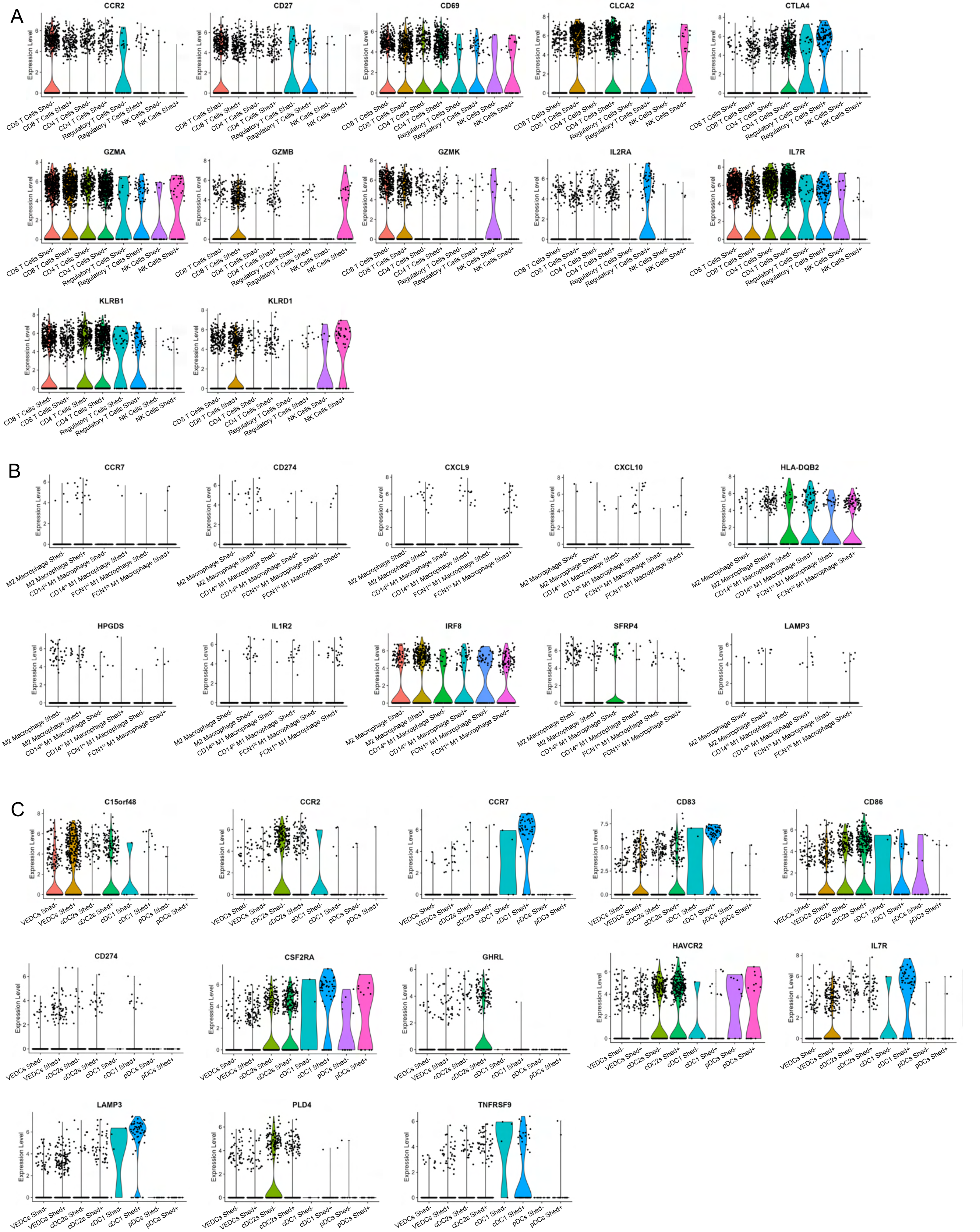

### Supplemental Figure 6

#### Supplemental Figure 6

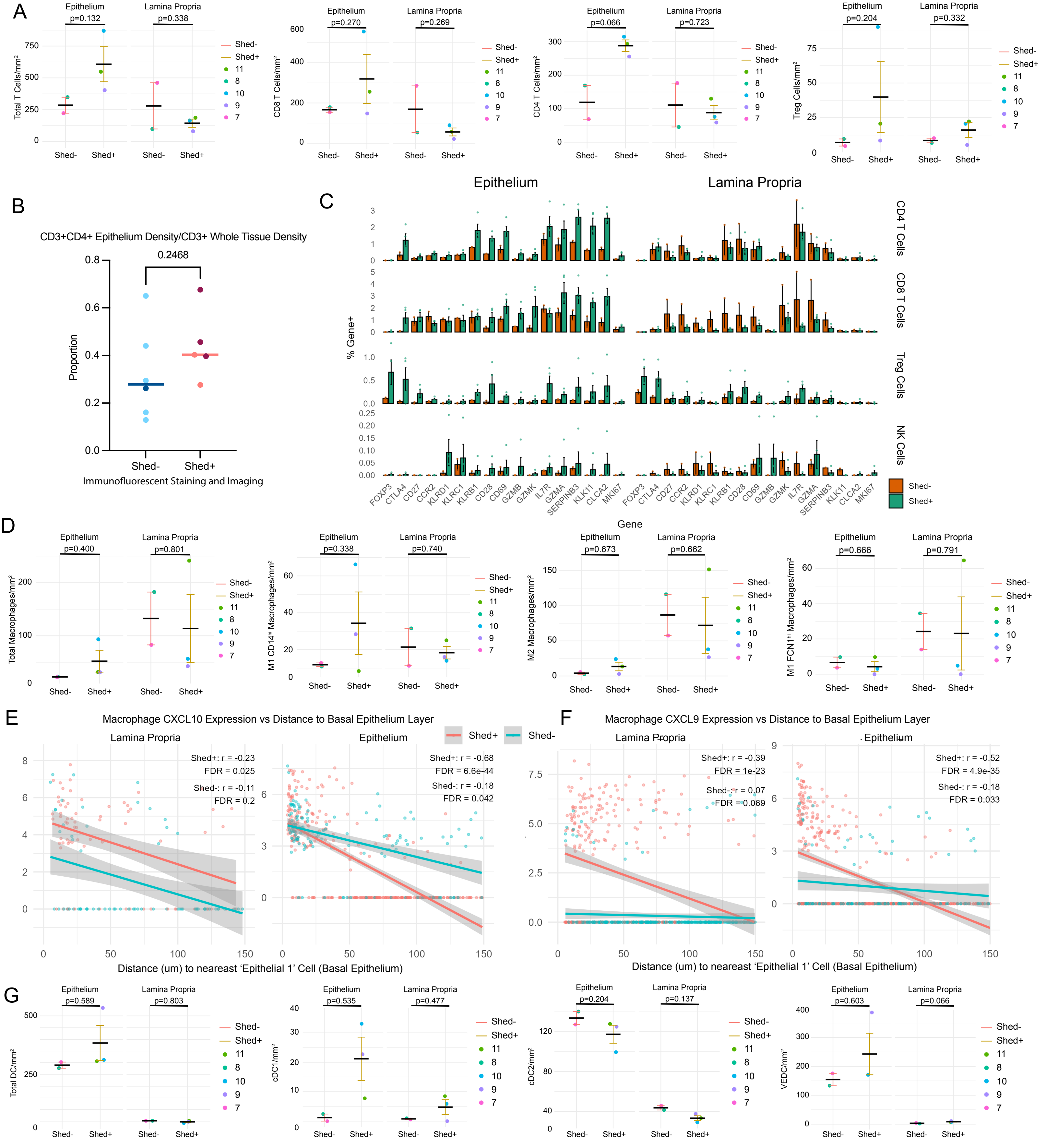

### Supplemental Figure 7

#### Supplemental Figure 7

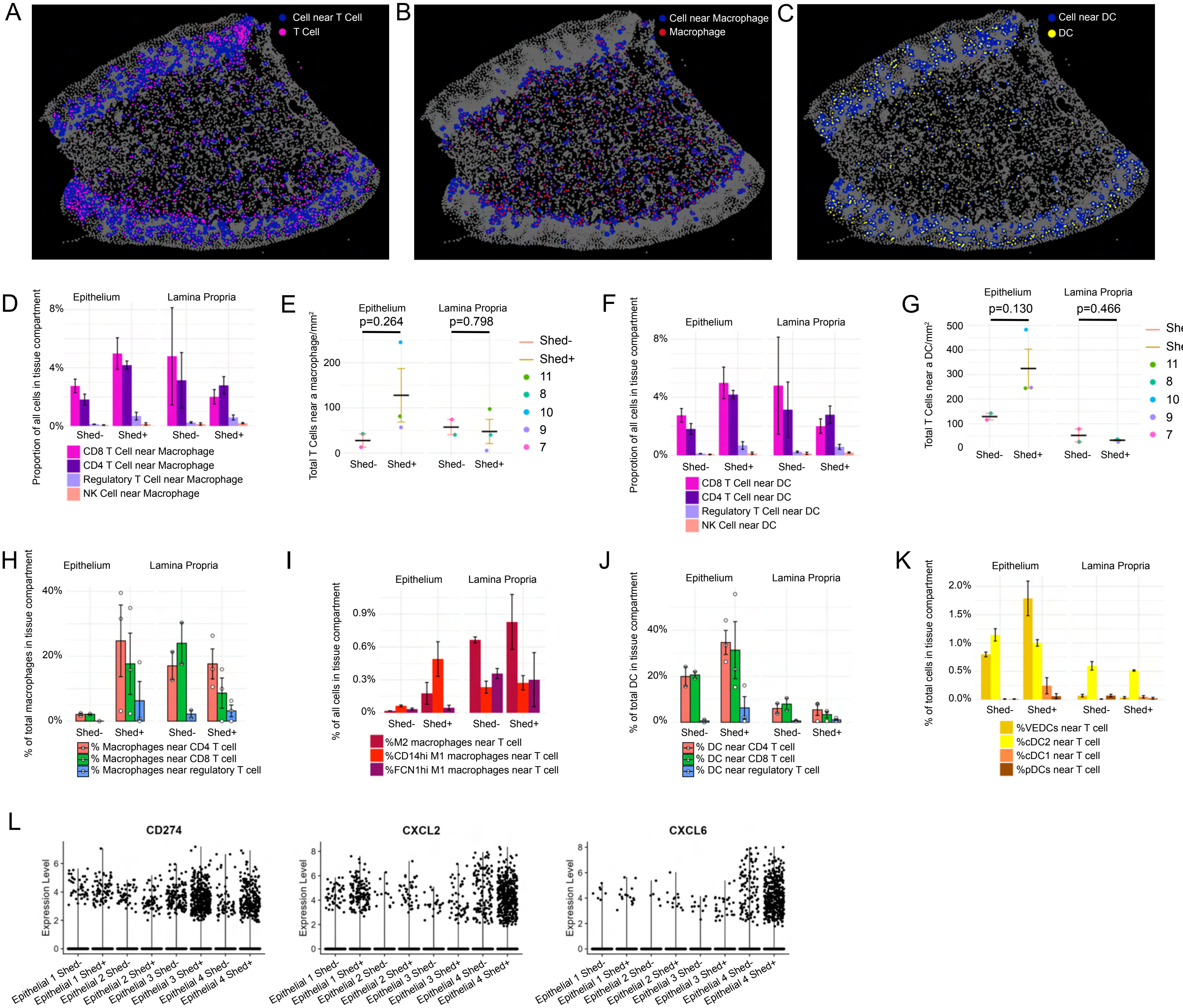
