## Supplemental Acknowledgments for "Genital herpes shedding episodes associate with alterations in the spatial organization and activation of mucosal immune cells"


**<u>Kinga Study Team:</u>**

*University of Washington: International Clinical Research Center*
Jairam R Lingappa (co-Principal Investigator and Protocol Chair), Justice Quame-Amaglo (study coordinator), Harald Haugen, Elena Rechkina (Laboratory Director), Daphne Hamilton, Matthew Ikuma, Marie Bauer, Zarna Marfatia, Kathy Thomas, Corinne Mar, Adino Tesfahun Tsegaye, Ayumi Saito

*Fred Hutchinson Cancer Center:*
Jennifer Lund (co-Principal Investigator), Sarah Vick, Finn MacLean, Jessica Graham, Jessica Swarts, Nicole Potchen, Irene Cruz Talavera, Lakshmi Warrier, Laura Pattacini, Paula Culver

*Kenya Medical Research Institute: Thika Partners in Health Research and Development (and Jomo Kenyatta University [JKUAT])*
Nelly R. Mugo (Site Principal Investigator), Kenneth Ngure (Site Investigator), Catherine Kiptinness (site coordinator), Bhavna H. Chohan (Site Laboratory Director), Nina Akelo, Stephen Gakuo, Elizabeth Irungu, Marion Kiguoya, Edith Kimani, Eric Koome, Solomon Maina, Linet Makena, Sarah Mbaire, Murugi Micheni, Peter Michira, Peter Mogere, Richard Momanyi, Edwin Mugo, Caroline Senoga, Mary Kibatha, Jelioth Muthoni, Euticus Mwangi, Philip Mwangi, Margaret Mwangi, Charles Mwangi, Stanley Mugambi Ndwiga, Peter Mwenda, Grace Ndung'u, Faith Njagi, Zakaria Njau, Irene Njeru, John Njoroge, Esther Njoroge, John Okumu, Lynda Oluoch, Judith Achieng Omungo, Snaida Ongachi, Dennis Wanyonyi, Everlyne Okong’o, Agnes Ndirangu
